# Neural effects of TMS trains on the human prefrontal cortex

**DOI:** 10.1101/2023.01.30.526374

**Authors:** Jessica M. Ross, Christopher C. Cline, Manjima Sarkar, Jade Truong, Corey J. Keller

## Abstract

How does a train of TMS pulses modify neural activity in humans? Despite adoption of repetitive TMS (rTMS) for the treatment of neuropsychiatric disorders, we still do not understand how rTMS changes the human brain. This limited understanding stems in part from a lack of methods for noninvasively measuring the neural effects of a single TMS train — a fundamental building block of treatment — as well as the cumulative effects of consecutive TMS trains. Gaining this understanding would provide foundational knowledge to guide the next generation of treatments. Here, to overcome this limitation, we developed methods to noninvasively measure causal and acute changes in cortical excitability and evaluated this neural response to single and sequential TMS trains. In 16 healthy adults, standard 10 Hz trains were applied to the dorsolateral prefrontal cortex (dlPFC) in a randomized, sham-controlled, event-related design and changes were assessed based on the TMS-evoked potential (TEP), a measure of cortical excitability. We hypothesized that single TMS trains would induce changes in the local TEP amplitude and that those changes would accumulate across sequential trains, but primary analyses did not indicate evidence in support of either of these hypotheses. Exploratory analyses demonstrated non-local neural changes in sensor and source space and local neural changes in phase and source space. Together these results suggest that single and sequential TMS trains may not be sufficient to modulate local cortical excitability indexed by typical TEP amplitude metrics but may cause neural changes that can be detected outside the stimulation area or using phase or source space metrics. This work should be contextualized as methods development for the monitoring of transient noninvasive neural changes during rTMS and contributes to a growing understanding of the neural effects of rTMS.

## Introduction

Repetitive transcranial magnetic stimulation (rTMS) is a safe and effective treatment for major depressive disorder, obsessive-compulsive disorder, smoking cessation, and migraines [1]. Despite FDA clearance for depression 15 years ago, response rates remain at 50% [2,3]. This suboptimal response rate may in part be because little is known about how rTMS treatment modulates neural activity in humans. Gaining a better understanding of how a single TMS train, the building block of rTMS treatment, modulates neural activity would provide foundational knowledge to guide the next generation of treatments. For example, development of an acute neural indicator of single TMS trains demonstrating prefrontal target engagement could guide high throughput screening of novel TMS patterns as well as adaptive, closed loop TMS treatments.

Unlike other noninvasive metrics, the TMS-evoked potential (TEP) provides a causal measurement of cortical excitability [4–11] and thus may be well suited for probing the neural effects of TMS trains. A few motor cortex studies have explored the acute neural effects of rTMS, demonstrating that a single TMS train modulates the evoked response within 55-100 ms of the train [12–14]. However, these results are difficult to interpret due to large TMS-related sensory responses evoked by each pulse in the train that overlap with the acute direct neural effects after the train [15–18]. Further, it is difficult to extend these motor cortex findings to the dlPFC where rTMS treatment is applied for depression. Indeed, little is known about the acute neural effects of TMS trains applied to the dlPFC, a region heavily implicated in psychiatric disorders [19]. In a prior study, we demonstrated that short-latency neural responses (25 and 33 ms after the train) may be larger following 10, 15, or 20 Hz trains compared to 1 Hz trains (manuscript under revision). While interesting, this study examined the evoked response to the *last pulse* in the TMS train (within 300 ms of the end of the train), rendering results difficult to interpret given the strong sensory responses from all pulses in the train lasting up to 300 ms after the last pulse in the train. To avoid these sensory confounds, in the current study we evaluated post-train effects at latencies greater than 300 ms after the last pulse in the train. To control for ongoing sources of variability in the brain, we used an evoked response to capture post-train excitability. To do this, we delivered two additional single TMS pulses starting at 500 ms after each TMS train and examined the resulting evoked responses.

For analysis of the post-train TMS-evoked potentials (TEPs), we focus on the early (< 80 ms) components of the TEP under the site of stimulation (early local TEP) as our primary outcome measure as it likely reflects local cortical excitability [20,21], tracks depression pathology [20], changes after rTMS treatment in depression, and these changes in a small study predicted clinical outcome [7,20]. A similar early evoked response is observed in invasive electrical recordings following single electrical pulses, providing some validation of the neural correlates of this component of the TEP [22]. Additionally, current TMS-EEG evidence suggests that the early TEP is less likely to be confounded by sensory responses [23–27].

In the current study, we sought to evaluate the acute neural effects of single and sequential dlPFC TMS trains using a sham-controlled, event-related study design. We hypothesized that single TMS trains but not sham would modulate acute cortical excitability, captured in the early local TEP, and that sequential TMS trains would lead to accumulated effects in the early local TEP. However, we did not observe single train or cumulative train effects on the early local TEP quantified using standard methods. In contrast, exploratory analyses revealed that single TMS trains may induce non-local effects on the evoked response as well as early local effects when using more sophisticated methods [28–30]. Together, this work provides foundational knowledge relevant for the monitoring of transient neural changes during rTMS that can be used to guide the next generation of treatments.

## Methods

### Participants

This study was carried out in accordance with the Declaration of Helsinki, reviewed and approved by the Stanford University Institutional Review Board, was performed in accordance with all relevant guidelines and regulations, and written informed consent was obtained from all participants. 52 healthy participants (19-65 years old [M=44.4, SD=13.3, 31F/20M/1O]) responded to an online recruitment ad and after an initial online screening and consent, 18 eligible participants (25-60 years old [M=42.9, SD=11.8, 9F/9M]) were enrolled. Two participants withdrew because rTMS was intolerable and the remaining 16 participants were included in the analyses (M=43.1 years, SD=12.5, 8F/8M). See Table S1 for additional demographics.

Inclusion criteria on the online screening form were (a) aged 18-65, (b) able to travel to study site, (c) fluent in English and (d) fully vaccinated against COVID-19. Exclusion criteria were (a) lifetime history of psychiatric or neurological disorder, (b) substance or alcohol abuse/dependence in the past month, (c) heart attack in the past 3 months, (d) pregnancy, (e) presence of any contraindications for rTMS, such as history of epileptic seizures or certain metal implants [31], or (f) psychotropic medications that increase risk of seizures including Bupropion (≥300mg/day) and/or any dose of Clozapine. Participants were also required to complete the *Quick Inventory of Depressive Symptomatology (16-item, QIDS)* self-report questionnaire and were excluded from the study if they scored 11 or higher indicating moderate depression [32,33]. All participants completed an MRI pre-examination screening form provided by the Richard M. Lucas Center for Imaging at Stanford University to ensure participant safety prior to entering the MRI scanner. Eligible participants were scheduled for two study visits: an anatomical MRI scan on the first visit and a TMS-EEG session on the second visit.

### Overall study design

Conditions were chosen to examine the transient neural effects induced after single and sequential 10 Hz TMS trains were applied to the left dlPFC (Fig 1A). We quantified transient induced neural effects using the TMS-evoked potential (TEP) evoked by single pulses of TMS (*probe pulses*) applied after each TMS train (Fig 1B). We chose to quantify the effects of TMS trains in this manner to obtain a causal measurement of train-induced neural effects and because TEPs are well described in the literature [7,34]. Because the sensory response to TMS pulses lasts for at least 300 ms [26,27,35–37], we chose to add a 500 ms delay between the last pulse in the TMS train and the first TMS *probe pulse*. Because we hypothesized that the neural effects of single TMS trains would be transient and not last longer than one second, and to examine the temporal specificity of the effects, we applied a second *control probe pulse* two seconds after the TMS train. While it would have been advantageous to probe the acute neural effects less than 500 ms after the TMS train, we determined that strong sensory responses to the TMS train and the inability to perfectly match the perception of active and sham TMS trains would render interpretation of a TEP < 500 ms after a TMS train difficult. For four subjects, we jittered probe latencies within early (500-700 ms) and later (1900-2100 ms) latency time windows to evaluate if the neural effects observed at 500 ms and 2 s were dependent on those exact timings (Fig S4). Given the lack of clear neural effects between TEP responses after probe pulses applied within each jittered 200 ms range (Fig S4), for subsequent subjects we focused experimentation on the neural effects at fixed probe times (500 ms and 2 s) following the TMS train.

**Figure 1.**
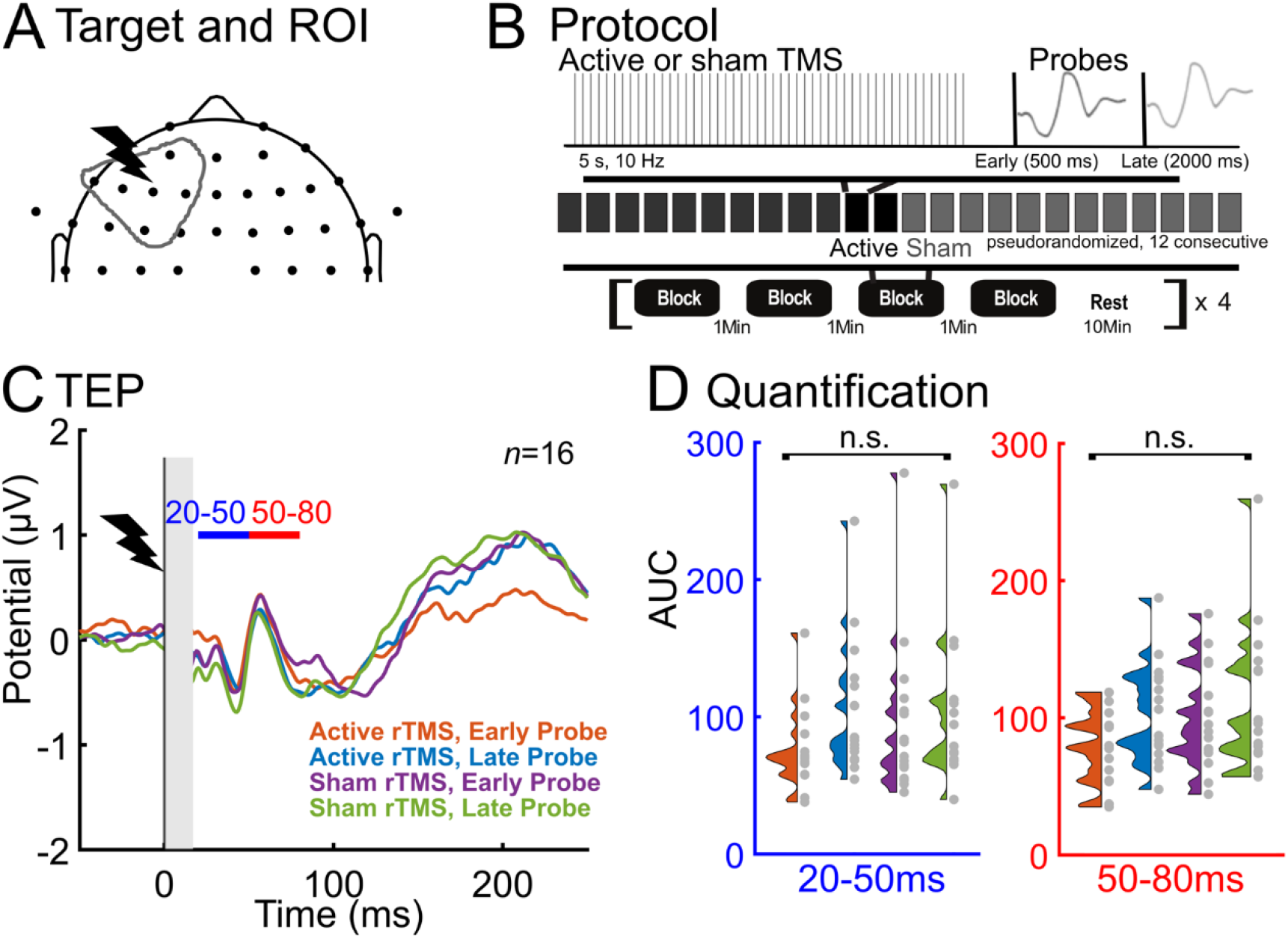
TMS trains did not modulate the sensor-space early local TEP amplitude. A) TMS was delivered over the left dlPFC and local TEP analysis was performed using six left frontal electrodes. B) Experimental design: Active and sham TMS trains were applied to the left dlPFC and single TMS pulses were used as probes to evaluate the TMS-evoked potential (TEP). Probe pulses were applied at 500ms (*early probe*) to assess early rTMS-induced changes or 2s (*late probe*) as a control. All probe pulses were active TMS, allowing a direct comparison between active and sham trains. To assess cumulative inter-train effects, up to 12 consecutive active and sham trains were applied. The blocks of active or sham rTMS were randomized. C) Group sensor-space TEPs (*n*=16) are shown for the -50ms to 250ms window. Time = 0 is the time of the single TMS probe pulse applied at 500 ms or 2 s after the TMS train. The two analysis latency windows are indicated with blue (20-50 ms) and red (50-80 ms) horizontal bars. See Supplementary for individual subject LMFA time series (Fig S1-S2). D) Null group effects of the local TEP size (quantified using LMFA) in both latency windows (20-50 ms, 50-80 ms) across conditions.

Our main outcome measure was the TEP from each probe pulse following TMS and sham trains. Changes in the TEPs were quantified by examining the early (20-50 ms and 50-80 ms) time windows of the TEP from EEG electrodes underneath the TMS brain target in the four conditions: (1) TEP from the probe pulse 500 ms after an active TMS train (*active rTMS, early probe*); (2) TEP from the probe pulse 2000 ms after an active TMS train (*active rTMS, late probe*); (3) TEP from the probe pulse 500 ms after a sham TMS train (*sham rTMS, early probe*); (4) TEP from the probe pulse 2000 ms after a sham TMS train (*sham rTMS, late probe*). To explore the cumulative effects of sequential TMS trains on the TEP, for each condition 12 consecutive TMS trains were applied in block order for 13 subjects. The order of conditions was randomized. We hypothesized that single 10 Hz trains to the dlPFC would affect the early local TEP for less than one second following active TMS trains (active rTMS early probe would be different than all other conditions). We further hypothesized that sequential TMS trains would enhance this effect.

### Transcranial magnetic stimulation

#### TMS targeting and calibration

Both single pulse TMS and TMS trains were delivered using a MagVenture Cool-B65 A/P figure-of-eight coil (MagVenture, Denmark) from a MagPro X100 system (MagVenture, Denmark). A thin (0.5 mm) foam pad was attached to the TMS coil to minimize electrode movement and bone-conducted auditory artifact. A TMS-Cobot-2 system (Axilum Robotics, France) was used to automatically maintain orientation of the active coil relative to the subject’s head. Neuronavigation (Localite TMS Navigator, Alpharetta, GA) was used to derive the TMS targets for each subject based on their individual T1-weighted MRI image. MRI was performed on a GE DISCOVERY MR750 3-T MR system (General Electric, Boston, Massachusetts) using a 32 channel head coil. T1 structural scans were acquired using a BRAVO pulse sequence (T1-weighted, sagittal slice thickness 1 mm, acquisition matrix 256 ✕ 256, TR 8 ms, TE 3 ms, FA 15°).

#### Resting motor threshold

To obtain resting motor threshold (RMT), single pulses of TMS were delivered to the hand region of the left primary motor cortex with the coil held tangentially to the scalp and at 45° from the midsagittal plane [38–40]. The optimal motor hotspot was defined as the coil position from which TMS produced the largest and most consistent visible twitch in a relaxed first dorsal interosseous (FDI) muscle [40]. RMT was determined to be the minimum intensity that elicited a visible twitch in relaxed FDI in ≥ 5/10 stimulations [41,42].

#### Determining target location, coil angle, and intensity

Our goal was to maximally modulate the left dlPFC node of the fronto-parietal network with TMS. Thus, we targeted a set of MNI coordinates (-38, 23, 38) previously identified as the group (*n*=38) peak of that node within the fronto-parietal network [43]. To minimize discomfort, we applied single pulses of TMS at 110% RMT at various angles (0°, 45°, and 90°) from the midsagittal plane and instructed subjects to select the angle that was most tolerable [27]. The optimal angle for each subject can be found in Table S2. Each subject then underwent a TMS train intensity ramp to introduce the sensation of the train. Subjects were instructed to notify operators if stimulation intensity became intolerable. The first TMS train of the visit began at 55% RMT and gradually increased to 110% RMT. In cases of intolerance, the stimulation intensity was adjusted down until tolerable for the following TMS train blocks (Table S2 for tolerable intensity used for each participant). Two participants found the TMS trains to be intolerable at all tested intensities and withdrew their participation. Eight participants could not tolerate the TMS trains at 110% RMT. To determine whether the size of the early local TEP was larger in subjects who received greater intensity rTMS, we performed a simple linear regression and found that TEP size was not significantly predicted by %MSO. To check whether the size of the early local TEP was larger in subjects with a greater cortical stimulation (accounting for individual factors such as anatomy and coil orientation), we calculated E-field magnitude for 14 subjects, using SimNIBS v3.2.5 [44,45]. Due to technical error during data collection we are not able to calculate E-field for 2 of our subjects. However, in the *n*=14 subjects we asked whether the size of the early local TEP was larger for subjects with a greater estimated E-field during rTMS, and found that TEP size was not significantly predicted by E-field (Fig S5 for more details).

#### Repetitive TMS (active rTMS)

We applied 10 Hz TMS trains for 5 s (50 pulses). These 5 s trains were used instead of 4 s trains following prior studies in an attempt to enhance the effects [46–49]. Each TMS train was followed by a first single pulse probe at 500 ms (*early probe*) and a second single pulse probe at 2000 ms (*late probe*), as defined in the previous section and depicted in Fig 1B. Probe TMS pulses were always ‘active’, regardless of whether they were preceded by an active or sham TMS train. Stimulation was arranged in 16 blocks. For *n*=3 subjects, trains were grouped within block in sequences of 3 or 6 sequential active or sham trains (in randomized order) for a total of 12 active and 12 sham trains per block. For the remaining *n*=13 subjects, trains were grouped in sequences of 12 active trains and 12 sham trains (in randomized order) per block so that cumulative effects of up to 12 sequential trains could be assessed. A 10-minute rest period was placed after every 4 blocks, and a 1-minute rest period between all other blocks.

#### Sham repetitive TMS (sham rTMS)

To quickly switch between active and sham TMS trains, we used a novel dual coil approach (Fig S13). The active TMS coil was positioned over the left dlPFC while the sham coil was positioned over the right dlPFC. The sham coil was a MagVenture Cool-B65 A/P coil with the sham side facing the scalp and fixed in place using a coil holder. Electrical current was delivered during sham TMS trains over the left frontalis muscle, using two surface electrodes (Ambu Neuroline 715) in order to approximate the somatosensory sensations arising from skin mechanoreceptors and scalp muscles during the active TMS condition [50]. We posited that although the sham TMS coil was placed over the right hemisphere, the left-sided electrical stimulation would be felt over the left hemisphere under the active TMS coil, the auditory click from the right-sided sham TMS coil would reach bilateral auditory cortices with similar timing and intensity as from the active TMS coil, and any perceived bone-conducted sound from the sham coil would not be precisely localized on the head. Although not an ideal setup, we felt it critical to be able to quickly switch between active and sham TMS and to our knowledge at present there is currently no commercially available approach for doing this. The intensity of the electrical current stimulation was calibrated to approximate the scalp sensation and discomfort of active TMS. To assess how closely matched the active and sham rTMS sensations were, subject perceptual ratings of loudness, scalp sensation, and pain were collected and analyzed (as described in more detail in Analyses).

### Electroencephalography

64-channel EEG was obtained using a BrainVision actiCHamp Plus amplifier, with ActiCAP slim active electrodes in an extended 10–20 system montage (actiCHamp, Brain Products GmbH, Munich, Germany) with a 25 kHz sampling rate to reduce the spread of the pulse artifact [51]. EEG data were online referenced to Cz and recorded using BrainVision Recorder software v1.24.0001 (Brain Products GmbH, Germany). Impedances were monitored and percentage of channels with impedances <10 kΩ was 94.38 ∓ 6.75%. Electrode locations were digitized using Localite (Localite TMS Navigator, Alpharetta, GA).

## Analyses

All EEG preprocessing and analyses were performed in MATLAB R2021a (Mathworks, Natick, MA, USA) using the EEGLAB v2021.1 toolbox [52] and custom scripts. TMS-EEG preprocessing was performed with the AARATEP pipeline, v2 [53], with source code available at github.com/chriscline/AARATEPPipeline. Data were processed in batches grouped by 4 sequential blocks (each batch containing probe responses from 48 active trains and 48 sham trains) to account for artifact changes that may occur over the duration of one session [54]. Epochs were extracted from 350 ms before to 1100 ms after each TMS probe pulse.

As part of the AARATEP pipeline, the following steps were taken, with all steps described in Cline *et al.* (2021) [53]. Data between 2 ms before to 12 ms after the pulse were replaced with values interpolated by autoregressive extrapolation and blending, downsampled to 1 kHz, and baseline-corrected using mean values between 350 to 10 ms before the pulse. Epochs were then high-pass filtered with a 1 Hz cutoff frequency and a modified filtering approach to reduce spread of artifact into baseline time periods (see [53] for details of this modified filtering approach). Bad channels were rejected via quantified noise thresholds and replaced with spatially interpolated values (see [53] for all details on channel deletion and interpolation); an average (+/- standard deviation) of 1.84 +/- 1.7 channels were rejected at this stage. Eye blink artifacts were removed using a dedicated round of independent-component analysis (ICA) and eye-specific component labeling and rejection using ICLabel [55]; an average (+/- standard deviation) of 1.75 +/- 0.55 components were rejected at this stage. Various other non-neuronal noise sources were attenuated with SOUND [56]. Decay artifacts were reduced via a specialized decay fitting and removal procedure (see [53] for details). Line noise was attenuated with a band-stop filter between 58-62 Hz. Additional artifacts were removed with a second stage of ICA and ICLabel component labeling and rejection, with rejection criteria targeted at removing any clearly non-neural signals (see [53] for all data deletion criteria); an average (+/-) standard deviation) of 64.9% +/- 7.31% of components, accounting for 41.0% +/- 21.6% of variance in the signal, were rejected at this stage. Data were again interpolated between -2 ms and 12 ms with autoregressive extrapolation and blending, low-pass filtered with a cutoff frequency of 100 Hz, and re-referenced to the average. We did not perform any trial rejection, since we found for this dataset that the AARATEP pipeline’s other processing steps (especially SOUND, decay fitting and removal, and multi-stage ICA rejection) produced sufficiently clean data.

### TMS train effects on the TEP

To compare evoked responses from TMS probe pulses at 500 and 2000 ms after active and sham TMS trains, we computed the local mean field amplitude (LMFA) for 20-50 ms and 50-80 ms time windows in a dlPFC region of interest (ROI). Because amplitude of an averaged EEG waveform is not independent of latency variance, and the early TEP peaks are not yet well defined, we used area under the curve (AUC), which captures the full morphology of the LMFA waveform by aggregating it over a time window rather than focusing on an instantaneous amplitude [57,58]. We chose the analysis time windows following Gogulski *et al.* (2023) [59] and because they are far enough after the interpolated time window (ending at 12 ms) and before time windows reported to include strong off-target sensory effects (>80 ms) [23,26,27]. The earlier 20-50 ms time window captures our primary hypothesized component of interest, the early local TEP. The later 50-80 ms time window captures other TEP components that have been previously reported in both prefrontal [59] and motor cortex [15–17]. The local ROI was chosen to cover the stimulation site and left lateral prefrontal cortex broadly: AF3, F7, F5, F3, F1, FC3. During the channel rejection procedure described above, an average (+/- standard deviation) of 0.62 +/- 0.65 channels and a maximum of 2 channels from this 6 channel ROI were rejected and spatially interpolated. TEP data were checked for standard assumptions using histograms and normal probability plots. Using a within subject design, the LMFA measurement from each time window (20-50 ms, 50-80 ms) was entered as a dependent variable into a repeated measures analysis of variance (ANOVA) with probe latency (early probe at 500 ms, late probe at 2000 ms) and stimulation (active TMS train, sham TMS train) factors. To test if TMS trains modulated neural activity in non-local regions, we repeated the above-described analysis using right frontal, left parietal, and right parietal ROIs. The parietal ROIs were selected because of relevance for fronto-parietal modulation by TMS when using our left dLPFC stimulation target [43].

### Sequential TMS train effects on the TEP

We examined the relationship between TMS train order (in the 12-train sequence) and TEP size following each train. To reduce dimensions, TEPs were averaged over groupings of three adjacent trains in the 12-train sequence (trains 1-3, 4-6, 7-9, and 10-12). Because there were 4 experimental blocks in this study, these groupings include 12 data points per group (3 trains × 4 blocks). To assess effects of TMS train sequence on the TEP, a repeated measures ANOVA was performed across these four train groupings. Three subjects did not have 12 sequential trains, so *n*=13 subjects were included in this analysis.

### Exploratory analyses

#### Sensor space hierarchical linear modeling (LIMO)

As an exploratory investigation of condition-specific TEP responses with minimal assumptions about relevant ROIs or time windows, we used the Hierarchical LInear MOdeling of ElectroEncephaloGraphic Data (LIMO EEG) toolbox [60]. LIMO is well-suited for exploratory investigations with minimal assumptions, which is why it is justified here. First-level beta parameters for each channel and time window were estimated from TEP features with ordinary least squares. Second-level analysis used a 2×2 repeated measures ANOVA (early probe, late probe; active rTMS, sham rTMS) with cluster-based correction for multiple comparisons (bootstrap *n*=1000, α = 0.05).

#### Phase space reconstructions using recurrence quantification analysis (RQA)

To evaluate whether TMS trains aligned the phase of neural oscillations (phase synchronization), we used recurrence quantification analysis (RQA). This is a dynamical systems approach to understanding complex dynamics that arise from phase shifting of oscillatory signals (see [30] for a review) and specifically for identifying transitions between coordinated and uncoordinated dynamics in a complex system [61]. In RQA, a time series is compared to itself with a predefined lag time to isolate phase regularity. Self-similarity of the time series is visualized using a recurrence plot, from which parameters are calculated to quantify aspects of phase regularity, such as predictability and complexity [28,29,62,63]. Signals that have high self-similarity (e.g. complex oscillatory signals) tend to revisit similar states and exhibit predictable dynamics. Patterns of predictable dynamics in a time series can be measured using the RQA parameter Percent Determinism [64,65]. Higher levels of determinism indicate coordination and stability (e.g. synchronization) of a system. This parameter was selected due to relevance for synchronization/desynchronization in neural circuit coordination and for distinguishing between coordination stability regimes [61,63,66]. Although less commonly used for TMS-EEG, we believe RQA to be justified for this investigation and may be more revealing about the nature of rTMS-induced phase dynamics than other methods for the following reasons: a) it is applied using a predefined lag which allows for focused study of specific EEG peaks, b) is used to understand phase dynamics agnostic to specific frequencies, and c) is valid for use with very short time windows compared with event-related spectral methods such as ERSP. For this analysis we used the local ROI and a 15 ms lag time to capture coordination dynamics relevant to early local TEP peaks such as N15-P30-N45. RQA was performed using the MATLAB RQAToolbox [28,29] (https://github.com/xkiwilabs) on EEG time series from 160 ms surrounding the evoked responses from early and late probe pulses after active and sham TMS trains (delay = 15 ms, embedding dimension = 6, range = -80 to 80 ms). Percent determinism was calculated using all trials and compared across the conditions with a 2×2 repeated measures ANOVA (active rTMS, sham rTMS; early probe, late probe).

#### Source space estimates

Subject-specific differences in gyral anatomy can cause underlying common cortical sources to project to the scalp in different topographies across subjects [67]. To account for this and other related consequences of EEG volume conduction, we performed EEG source estimation. Using digitized electrode locations and individual head models constructed from subjects’ anatomical MRI data [45], subject-specific forward models of signal propagation from dipoles distributed over and oriented perpendicular to the cortical surface to electrodes on the scalp were constructed [68,69]. One subject did not have digitized electrode locations available, so was excluded from the source analysis (*n*=15). Inverse kernels mapping the measured scalp EEG activity to underlying cortical sources were estimated using weighted minimum-norm estimation (wMNE) as implemented in Brainstorm [70]. Surface-based morphometry was used to map activations from subject-specific cortical surfaces to a common group template surface (ICBM152). A data-driven process was used to identify source-space spatial filters based on observed peaks in the average source TEP responses aggregated from data pooled across all subjects and stimulation conditions. Response amplitudes were then extracted by applying these latency-specific spatial filters to subject-and condition-specific data subsets. This approach to analyzing EEG in source space without prior assumptions about time window or connectivity is justified here to support our exploratory goals and to guide future investigations. For each identified TEP latency of interest, a 2×2 repeated measures ANOVA was used to assess effects of stimulation (active rTMS, sham rTMS) and probe latency (early probe, late probe).

### Sensory perception of TMS trains and single pulse TMS probes

To assess how closely matched the active and sham conditions were in sensory perception, subjects rated their perception of loudness, scalp sensation, and pain. Participants were instructed to respond verbally immediately following each stimulation to rate *loudness, scalp sensation,* and *pain perception* on scales ranging from 0 to 10. To ensure consistency in how these questions were phrased across conditions and subjects, scripts were used following Ross *et al.* (2022) [27]. In each study visit and prior to the experimental conditions, participants were asked to provide perceptual ratings after each of seven stimulation conditions: (1) single pulse TMS with no preceding train, (2) active TMS train with no probe pulse, (3) early probe at 500 ms after an active TMS train (4) late probe at 2000 ms after an active TMS train, (5) sham TMS train with no probe pulse, (6) early probe at 500 ms after a sham TMS train, and (7) late probe at 2000 ms after a sham TMS train. The order of these conditions was randomized across participants but with single pulse TMS perceptual testing applied before all TMS train conditions for all participants. This order of single pulse perceptual ratings followed by TMS train perceptual ratings was performed to avoid possible carry-over of the feeling of the TMS train affecting the perceptual score of the single TMS pulse.

#### Statistical analyses of perceptual ratings

Perceptual ratings were compared between active and sham conditions (active vs. sham rTMS, early probe after active rTMS vs. early probe after sham rTMS, late probe after active rTMS vs. late probe after sham rTMS) using nine paired t-tests for loudness perception, scalp feeling, and pain ratings. Multiple comparisons were corrected for using the Bonferroni type adjustment.

## Results

### TMS train effects on the TEP

First we asked whether there was a significant change in the local sensor-space TEP amplitude after different types of TMS trains (active vs. sham) and using different post-train probe latencies (500 ms, 2000 ms). This analysis revealed that single trains of left dlPFC TMS did not affect the amplitude of the TEP based on typical response metrics (Fig 1, *n*=16). When analyzing the first peak in the early local TEP (20-50 ms, LMFA, Fig 1D and S1-S2), we found no effect of stimulation (F(1,15)=0.9735, p=0.3395) but a main effect of probe latency (F(1,15)=10.2000, p=0.0060), and no stimulation by probe latency interaction (F(1,15)=3.1384, p=0.1000). To assess whether this null stimulation effect was due to the specific latency chosen between the TMS train and probe pulses, four subjects with jittered early probe latencies (500-700ms) were evaluated; we found no significant effect on TEP response (see Fig S4 for more details). To verify that the null results of TMS trains on the early local TEP were not due to the type of quantification performed (LMFA), we repeated the analysis using peak to peak amplitudes (in dB) and observed no main effect of stimulation (F(1,9)=0.9619, p=0.3523; see Fig S14) and no main effect of probe latency (F(1,9)=0.5255, p=0.4869). We found similar results when analyzing the later peak in the early local TEP (50-80 ms, Fig 1D and S1-S2); we observed no effect of stimulation (F(1,15)=2.2449, p=0.1548) and no effect of probe latency (F(1,15)=4.2198, p=0.0578). We found similar results when analyzing a wider time window in the early local TEP (20-80 ms); we observed no effect of stimulation (F(1,15)=2.1422, p=0.1639) but a main effect of probe latency (F(1,15)=7.4435, p=0.0156), and no stimulation by probe latency interaction (F(1,15)=3.6761, p=0.0744). We also analyzed the N100 of the TEP (80-130 ms) and observed no effect of stimulation (F(1,15)=2.2449, p=0.1548) and no effect of probe latency (F(1,15)=4.2198, p=0.0578). We also analyzed the P200 of the TEP (130-250 ms) and observed no effect of stimulation (F(1,15)=0.9735, p=0.3395) but we did find a main effect of probe latency (F(1,15)=10.2003, p=0.0060). We found no stimulation by probe latency interaction (F(1,15)=3.1384, p=0.0968). See Figs S1-S2 for individual subject TEP waveforms across conditions. In summary, active TMS trains did not differ from sham TMS trains in eliciting early TEP sensor-space amplitude effects at the site of stimulation.

We next investigated the TEPs from non-local ROIs, at regions expected to be connected to the stimulation site, and quantified the effects of stimulation type (active, sham TMS trains) and probe latency (early, late probes). To do so, we calculated the global mean field amplitude (GMFA, using all electrodes) and local mean field amplitude (LMFA) in regions of interest (ROIs) farther from the site of stimulation (right frontal, left parietal, right parietal; Fig 2, *n*=16, see Methods for more details). GMFA and LMFA of the ROIs appear to have early TEP peaks in a 20-50 ms time window. Using all electrodes (GMFA), we observed in this time window no main effect of stimulation (Fig 2C, E; active vs. sham TMS trains; F(1,15)=3.1534, p=0.0961) but an effect of probe latency (Fig 2C, E; early probe vs. late probe; F(1,15)=7.2050, p=0.0170), and no stimulation by probe latency interaction (F(1,15)=4.2375, p=0.0573). In the right frontal ROI, we observed a main effect of stimulation (F(1,15)=6.0103, p=0.0270) such that active trains reduced TEP size compared with sham trains, a main effect of probe latency (F(1,15)=7.8531, p=0.0134) such that the TEPs from the early probe were reduced compared with the TEPs from the late probe, and no interaction between stimulation and probe latency (F(1,15)=1.4014, p=0.2549). In the other two ROIs, there were no main effects of stimulation: in the left parietal ROI, we found no effect of stimulation (F(1,15)=4.4061, p=0.0531) and no effect of probe latency (F(1,15)=2.1164, p=0.1663) and in the right parietal ROI, we found no effect of stimulation (F(1,15)=0.7821, p=0.3904), an effect of probe latency (F(1,15)=5.3080, p=0.0360), but no interaction (F(1,15)=2.3411, p=0.1468). See Fig 2D,F for more details, and supplementary for individual subject GMFA (Fig S6) and non-local LMFA time series (Figs S7-S9), including in the 20-50 ms time window and also in a 50-80 ms time window. In summary, we observed that in the right frontal ROI there was a main stimulation effect of TMS train type that represents a reduction in TEP size with active TMS trains, a main effect of probe latency that represent that the TEP size reduction is no longer present by 2000 ms after the train, and no clear effects in GMFA or other non-local ROIs tested in either of the early TEP time windows.

**Figure 2.**
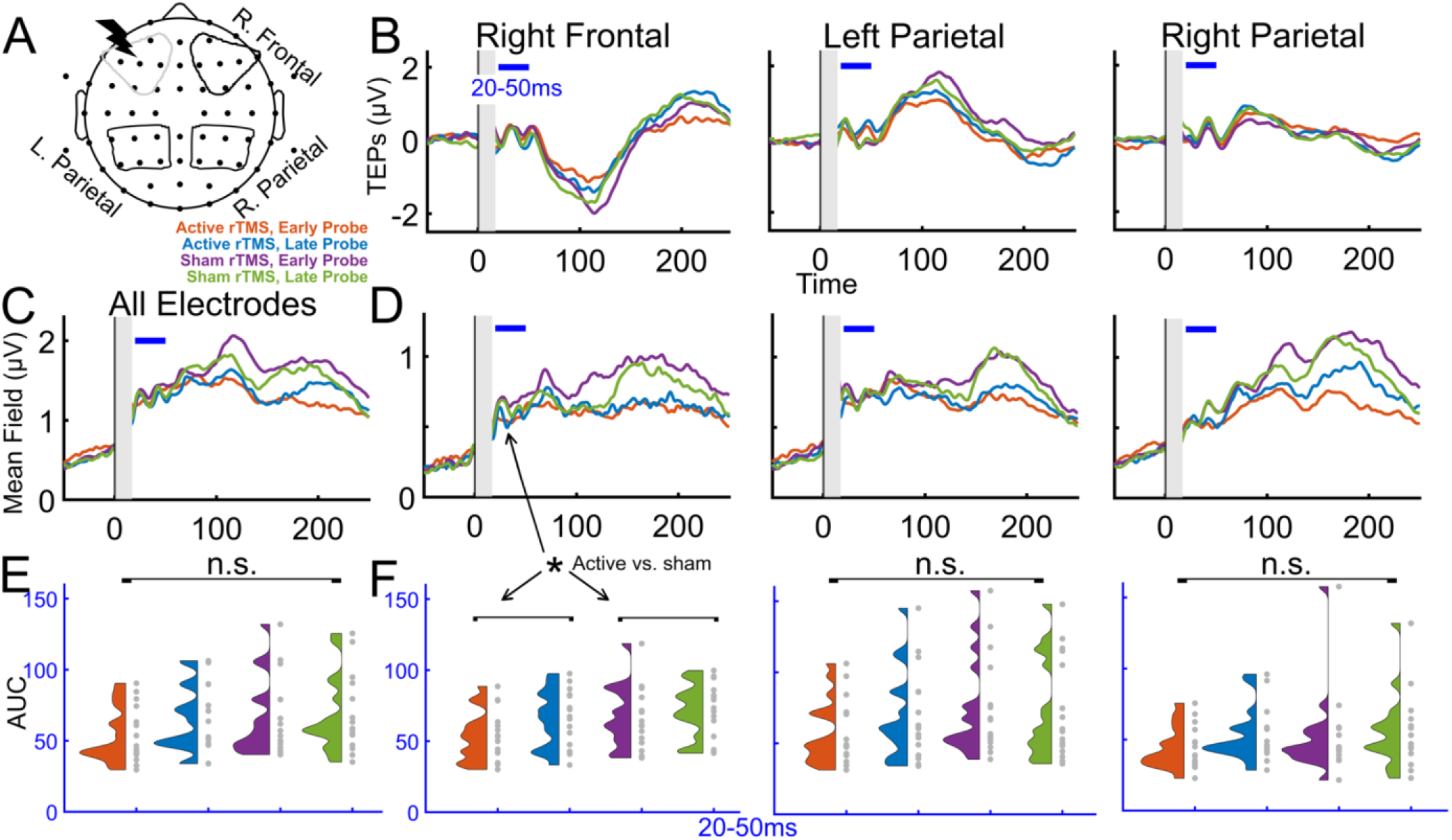
TMS trains reduce cortical excitability in the right frontal cortex. A) Regions of interest for non-local TEP analyses. B-F) Comparisons of TEP conditions using all electrodes (GMFA) and downstream ROIs (LMFA). B) Group TEPs (*n*=16) for each ROI, with the 20-50 ms analysis time window indicated with a blue horizontal bar. C) Group averaged global mean field amplitude for all electrodes (*n*=16). D) Group average local mean field amplitude for non-local ROIs. E) Individual subject global mean field amplitude area under the curves in the four conditions. F) Group LMFA across conditions. See Supplementary S6-S9 for individual subject time series and results from both 20-50 ms and 50-80 ms time windows. The main effect of stimulation is indicated with an asterisk (*p<0.05).

### Sequential TMS train effects on the TEP

Next, we asked if repeated TMS trains elicited cumulative neural effects, assessed by sequential TEP measurements (Fig 3, *n*=13). We observed no cumulative effects on the early local TEP after up to 12 sequential TMS trains. See Figure 3B-C (20-50 ms window; F(3,36)=0.4340, p=0.7300) and Figures 3B and S10 for the 50-80 ms window (F(3,36)=0.7722, p=0.5172). For an additional follow-up analysis of later TEP time windows 80-130 ms (Fig S11; F(3,36)=2.6313, p=0.0648) and 130-250 ms (Fig S12; F(3,36)=1.2487, p=0.3065), see Supplementary Materials. In summary, we did not observe a group effect of sequential TMS trains on the TEP.

**Figure 3.**
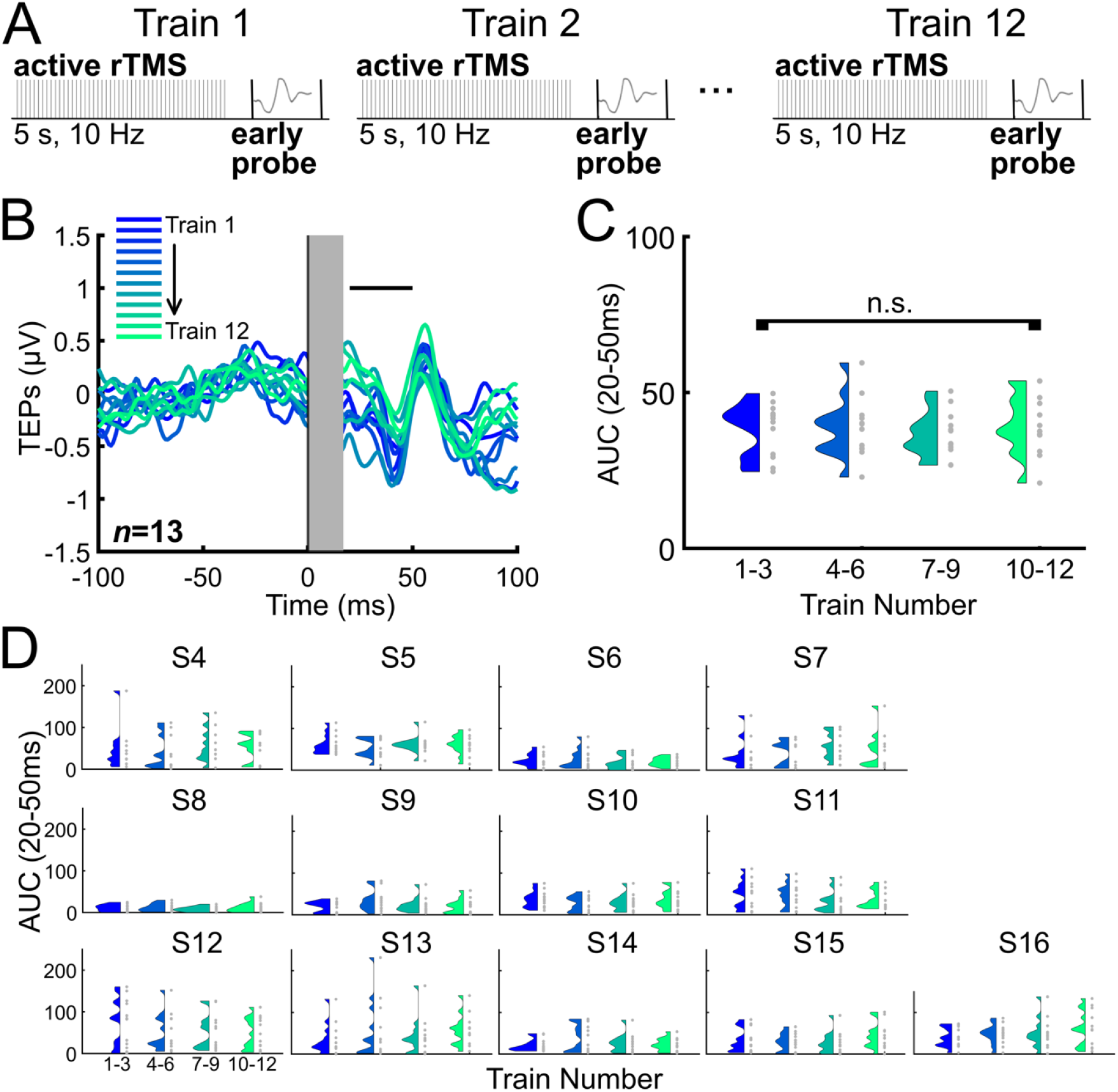
Sequential TMS trains did not modulate the early local TEP. A) 12 active TMS trains were applied sequentially. Each of the 12 trains were followed by an early TMS single pulse probe at 500 ms (also see Fig 1B). B) Group average (*n*=13) TEPs from the early probe following each sequential active TMS train, with the early latency window indicated with a black (20-50 ms) horizontal bar. C) Group average TEP response from the early probe following each sequential active TMS train. TEPs are grouped based on the order of the associated TMS train (groupings of three consecutive trains). D) Individual subject TEP results from the early probe following sequential active TMS trains. For the 50-80 ms TEP window see Fig S10. For later latencies see Figs S11-S12.

### Exploratory analyses

#### Sensor space hierarchical linear modeling (LIMO)

As an exploratory investigation of condition-specific responses with minimal assumptions about relevant ROIs or time windows, we used the LIMO toolbox [60]. LIMO analyses revealed significant effects of stimulation (active vs. sham), of probe latency (early vs. late), and significant interactions between these two factors. The most prominent effects were observed at later latencies of the TEP between 100-250 ms at central and bilateral frontal scalp electrodes (Fig 4A, *n*=15).

**Figure 4.**
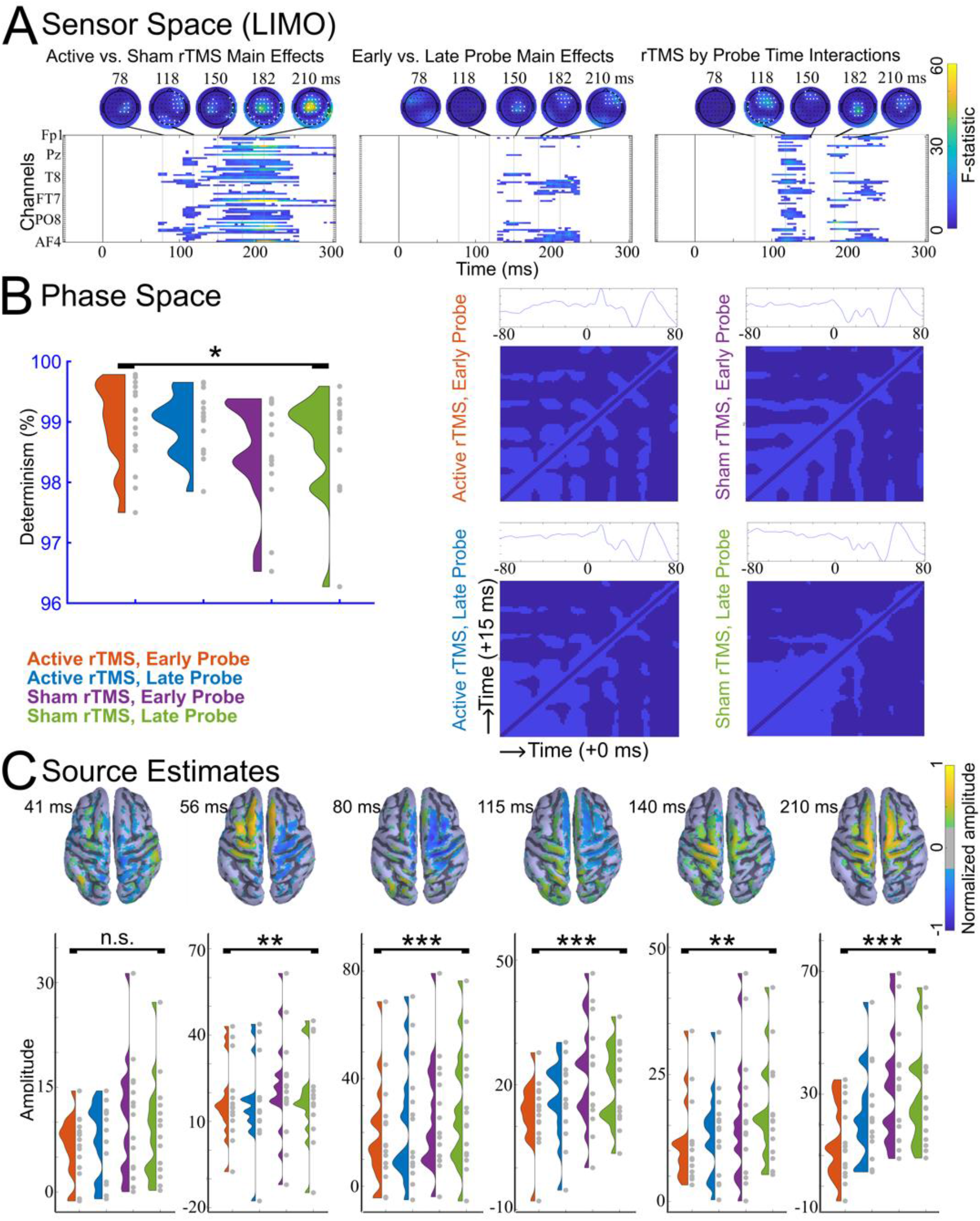
Exploratory analyses revealed that TMS trains modulated the non-local late TEP, resulted in more predictable phase dynamics in the early local TEP, and reduced the size of source estimates across both early and late TEP components. A) Sensor space hierarchical linear modeling analysis (LIMO; *n*=16) F-statistic heatmaps and topographies, corrected for multiple comparisons. Electrodes with F-statistics surpassing the significance threshold are shown in white. B) Group averages of percent determinism by condition and RQA recurrence plots averaged over all subjects (*n*=16; range in ms on each axis, lag = 15 ms, embedding dimensions = 6). C) Source estimates for condition specific TEPs (*n*=15) with topographies and group averages shown at 6 peak times that were determined from the source TEP averaged across all four conditions. For more details, see Fig S15. Main effects of stimulation are indicated with asterisks (*p<0.05, **p<0.01, ***p<0.001).

#### Phase space reconstructions using recurrence quantification analysis (RQA)

EEG complexity due to oscillatory phase shifting was quantified using RQA [28–30]. This analysis on percent determinism revealed significant main effects of stimulation (F(1,15) = 5.2452, p = 0.0369), no main effects of probe latency (F(1,15) = 0.3869, p = 0.5433), and no stimulation by probe latency interactions (F(1,15) = 0.4596, p = 0.5082). Percent determinism was greater following active than sham TMS trains, regardless of probe latency (Fig 4B, *n*=16). For a few subjects, percent determinism appeared lower than the group (96.27, 96.84, 96.53, 97.50). This metric is unlikely to have true outliers due to artifactual or spurious data and the qualitatively high percentage in all subjects (>96% DET) is indicative of normal brain EEG [71,72]. However, due to the susceptibility of a small sample to individual subject data points, we verified that none of these data points were outside of 3 standard deviations from the mean (2.87, 1.93, 2.28, and 2.12 standard deviations below the mean).

#### Source space estimates

To extract data-driven latency-specific source ROIs, cortical source activity was averaged across all stimulation conditions and subjects (Fig 4C, top row). 2×2 repeated measures ANOVAs were performed on amplitudes at eight peak times that were identified in the average source response time series. We found smaller source space TEP peaks following active TMS train conditions compared to sham TMS train conditions, at latencies spanning 56-210 ms. Specifically, we report main effects of stimulation type on source space amplitudes such that source TEPs are smaller following active TMS trains than following sham TMS trains at 56 ms (F(1,14)=12.077, p=0.00371), 80 ms (F(1,14)=21.576, p=0.000379), 115 ms (F(1,14)=17.442, p=0.000932), 140 ms (F(1,14)=14.045, p=0.00216), and 210 ms (F(1,14)=39.473, p=0.0000201). We also found a main effect of probe latency on the source space TEPs; at 140 ms source space TEPs were smaller when probed at 500 ms than when probed at 2000 ms (main effect of probe latency at 140 ms, F(1,14)=4.837, p=0.0452). We found two stimulation by probe latency interactions later in the source space TEP, at 115 ms (F(1,14)=13.502, p=0.00250) and at 210 ms (F(1,14)=18.302, p=0.000765). See Fig 4C for all group level source space estimate time series and topographies, Fig S15 for more details, and Fig S16 for all individual subject source space TEPs. In summary, we found that there may be active TMS train-induced source-space TEP reductions across the TEP latencies, and that at a 115 ms TEP peak this effect was greater for earlier probes compared to late probes (Fig 4C).

### Sensory perception of TMS trains and single pulse TMS probes

To better understand how well matched the sensory experiences were between active and sham TMS train conditions, we compared the perceptual ratings (Fig S3) using paired t-tests. These nine tests were corrected for multiple comparisons, resulting in an adjusted α of 0.0056. With respect to isolated active vs. sham TMS trains, we observed no statistical differences in loudness perception (t(15) = 3.8776, p = 0.0015), pain (t(15) = 1.0274, p = 0.3205), or scalp sensation (t(15) = 3.0138, p = 0.0087). When testing the perception of the probe pulses following active vs. sham TMS trains, we observed no effects across all conditions, including loudness perception (early probe, t(15) = 2.6112, p = 0.0197; late probe, (t(15) = 2.3342, p = 0.0339), scalp sensation (early probe, t(15) = 3.0138, p = 0.0186; late probe, t(15) = 2.893, p = 0.112), and pain (early probe, t(15) = 2.9084, p = 0.0108; late probe, t(15) = 3.0382, p = 0.0083)). The only comparison that resulted in statistically significant differences was loudness perception between active and sham TMS trains (t(15) = 3.8776, p = 0.0015), indicating that the active TMS trains were perceived to be louder than the sham trains. These results suggest that the perception of scalp feeling and pain were relatively well matched between active and sham conditions, that auditory loudness perception was indistinguishable across different probe latencies within train conditions, but that loudness perception was unmatched between active and sham TMS trains.

## Discussion

In this study we sought to evaluate whether 10 Hz TMS trains to the dlPFC induce acute neural effects quantifiable in the early (20-80 ms) local TMS-evoked potential (TEP). Due to the well-described sensory responses to TMS, which we also observed after each pulse within a train, we applied the probe single pulses at latencies that should be clear of train-evoked sensory potentials (>300 ms). We found that: 1) prefrontal TMS trains did not induce acute neural changes in the size of the sensor-space early TEP in regions local to the stimulation site but 2) suppressed the early TEP in a right frontal ROI when probed at 500 ms, and that 3) up to 12 sequential TMS trains did not elicit a cumulative effect on early local TEPs. In exploratory analyses, we found that TMS trains may induce 1) neural changes non-locally at later latencies (>78 ms), 2) regularity in early local oscillatory phase dynamics, and 3) reduced source estimate amplitudes spanning 56-210 ms. In the context of previous work [12,13,53], these findings indicate that TMS trains did not strongly modulate neural excitability in the left prefrontal cortex and exploratory analyses have implications for the direction of future work.

### Sequential TMS train effects on the TEP

There are many open questions about the relationship between the amount of TMS (number of pulses, trains, or sessions) and clinical outcomes [2,73–75]. This discussion about accumulated effects of rTMS is also relevant for neurophysiological outcomes, such as the motor-evoked potential (MEP [76–78]) and EEG [77]. In this study we asked whether accumulated neurophysiological effects of rTMS could be measured across 12 rTMS trains. Previous work demonstrated that 12 trains of 5 Hz rTMS has a larger effect on MEP amplitude than 1 train [79], but how this finding translates to EEG responses measured after 10 Hz stimulation is unknown. However, because it is generally accepted that when enough TMS trains are applied to the dlPFC there are lasting neural changes, we hypothesized that sequential TMS trains would induce changes in cortical excitability that accumulate as a function of TMS train. We observed that up to 12 sequential TMS trains was not sufficient to induce cumulative neural changes in the early TEP measured locally to the site of stimulation. Several possible explanations exist: 1) 12 trains represents only ∼1/6 of a standard TMS treatment for depression and thus is not sufficient to modulate cortical excitability; 2) 12 trains is sufficient but the resulting changes in cortical excitability are not best captured using an ANOVA and TEP sensor space amplitudes; 3) low signal-to-noise in the TEP metric reduced the ability to detect change; 4) 12 trains is sufficient in depressed but not in healthy participants. Regardless of the reason, future work is needed to tease apart this critical question. We are developing a better understanding of how the signal-to-noise of the TEP differs as a function of dlPFC subarea [59] and are now developing fully personalized methods to minimize artifact and boost signal. We are also developing methods to better match the perceptual effects of active and sham TMS trains so that small neural changes in active compared to sham rTMS are more easily detected and attributed to direct effects and not differences in sensory perception. Future work will continue to investigate this important question about neural changes from sequential TMS trains by applying more sequential TMS trains. It is worth noting that experimental limitations could lead to a notable trade-off between longer sequences of TMS trains and a well-controlled, randomized study design: more sequential TMS trains would require greater study resources for similar sham matching.

### Non-local effects of TMS trains

We observed that TMS trains can modulate cortical excitability in non-local brain regions. Specifically, 10 Hz trains modulated cortical excitability in the right frontal cortex (Fig 2B-F). Responses in EEG from propagation to the right frontal cortex within this time window have been reported previously with TMS to left dlPFC [80] and cerebellum [81] and could reflect network effects of TMS in the hemisphere contralateral to our dlPFC target. We also analyzed the TEPs with LIMO, which is particularly well-suited for exploratory investigations of EEG across spatial and temporal scales [60]. The strength of this approach is for hypothesis generation starting with minimal assumptions about relevant ROIs or time windows. TMS appeared to reduce cortical excitability at the scalp vertex and along the midline in later TEP latencies (>78 ms). Considering the suggested role of inhibitory influences in this later time window [18,26], these results could suggest that our TMS protocol reduced inhibitory contributions to the TEP. However, due to the known sensory-related vertex potential that occurs within this same time window [23,26,27], these results may also be interpreted to indicate perceptual differences between the experimental conditions. Our further exploratory analysis in source space supports that active TMS may reduce neural activity at multiple latencies (spanning 56-210 ms) after the TMS probe pulse. However, these reductions were not generally localized to the site of stimulation. There are two possible explanations for these non-local results. One is that these non-local results reflect vertex potentials due to condition-specific sensory effects of TMS trains. The other possible explanation is that the non-local changes are from indirect, network-level effects. For example, if TEP peaks in this time window are inhibitory in nature as some have suggested [82,83], our data would support that TMS trains may reduce inhibition in connected brain regions. However, as this was an exploratory analysis we did not have any predictions for modulation in these regions.

### Early local phase dynamics resulting from TMS trains

To address the possibility that the TEP is not suited for capturing rTMS-induced phase coordination behavior, we applied a dynamical systems method for quantifying changes in phase space of time series that consist of oscillatory signals, RQA [28]. This method was used to capture neural coordination behavior (synchronization/desynchronization) modulated by 10 Hz TMS trains. One strength of RQA is that it can robustly measure complexity of a signal that is short duration and non-stationary, as in short recordings of human EEG [63,71,72,84–87]. We found that active TMS trains can have quantifiable effects on EEG phase dynamics – that the phase of the early local TEP may have more determinism following active TMS trains than following sham trains. If this result replicates in future work, it would indicate increased predictability following active TMS trains. Because high determinism can be indicative of a stable regime within metastable or multistable coordination dynamics, as in multifunctional neural coordination [66], increased determinism in the TEP would likely represent phase synchronization. This analysis indicates that TMS trains may induce predictable phase dynamics in TMS-affected neuronal populations [28,66]. This dynamical systems approach may be valuable in future investigations to help us understand how rTMS modulates neural coordination [66], which is not possible from quantifying TEP amplitude alone.

### Limitations and future directions

One important limitation of the work is the small sample size. The small sample limits our ability to make statistically robust and generalizable conclusions from the work. These findings should be tested in a larger sample of subjects for statistical robustness and generalizability. Additionally in future work, any robust findings should also be examined at the individual subject level to determine relevance for individualized patient treatments.

Other possibilities for the null effects from single or multiple trains on TEP amplitudes are related to our methodological decisions. First, the exact probe latencies of 500 and 2000 ms may not have been ideal to capture neural effects. In a subsample of subjects, we found that jittering the early and late probe latencies from 500-700 ms and from 1900-2100 ms, respectively, did not influence the early local TEP amplitude (Fig S4). Secondly, it is possible that the acute changes from the TMS train resolve by 500 ms and thus the TMS probe pulse was not applied close enough to the train to capture these effects.

Unfortunately, applying probes earlier than 500 ms would likely result in TEPs that are confounded by sensory artifacts from the pulses within the train. Unless active and sham TMS trains are perfectly matched perceptually, the neural effects when placing the early probe closer to the TMS train would be difficult to interpret. It is worth noting that the timing of the single probe pulse was placed outside of the range of the sensory effects of the TMS train for two reasons: 1) because of the lack of perfectly perceptually matched sham train control, and 2) with an eye towards exploring the neural effects of other train frequencies, amplitudes, and patterns. Even if sham can be perfectly matched perceptually, the train differences between active conditions varying in frequency, intensity, or patterns will undoubtedly result in different perceptual effects. Hence, we feel that too many confounds exist between 0-300 ms after the TMS train to probe neural effects of perceptually un-matched train parameters using existing methods. Thirdly, rTMS dosing and coil orientation both were adjusted in some participants due to tolerability issues. These parameters were held constant within subject across all conditions, critically important for the within-subjects design. However, it is known that these parameters are related to the magnitude of brain response and to which neurons are most affected [88], and thus it is possible that the variation in these parameters could result in some subjects having larger rTMS effects across all active conditions than other subjects. Using regression analyses we found that the size of the early local TEP was not significantly larger for subjects with a greater %MSO or estimated E-field during rTMS, but additional studies are needed with larger sample sizes to fully address the role of individualized rTMS parameters in the strength of TMS effects.

Although our primary results were null, exploratory analyses indicate that the study design may be sufficient to capture changes in the TEP in non-local brain regions at later latencies, in phase coordination dynamics, and in distributed source estimates. These analyses provide the basis for motivating hypotheses for future testing that train effects may be examined as synchronization of excitatory and inhibitory neural activity in prefrontal cortex and connected networks. Future work should explore the EEG signal as a dynamical system with phase shifting behavior, and as such to characterize changes in this behavior following TMS trains to better understand how TMS modulates complex neuronal population synchronization. Phase coordination of EEG may provide the basis for a more detailed understanding of TMS-induced brain changes than by quantifying changes in the size of the TEP. Ongoing work in our lab will further explore neural complexity that may arise from application of TMS trains. As mentioned above, this train-probe approach described here can also be used to optimize the neural effects of different TMS train parameters for the next generation of treatment protocols. Finally, future work should explore whether this experimental approach can be used for real-time measurement and monitoring of the neural effects of TMS trains. If successful, this real-time approach would have important implications in developing closed-loop adaptive TMS treatments [89].

Finally, given our choice of stimulation parameters (5 s 10Hz train) is different from the FDA-cleared 10 Hz protocol (4 s train) and the recently cleared theta burst stimulation (TBS) protocol, it is possible that stimulation with these other parameters may elicit different neural effects. Thus, our choice of stimulation parameters in this study limits generalizability. To better understand the intricate interplay between stimulation parameters and acute neural effects, future investigations need to systematically explore the neural effect of other stimulation parameters.

## Conclusions

We evaluated the neural effects of single and sequential 10 Hz TMS trains to the left prefrontal cortex using single ‘probe’ TMS pulses. Single left hemisphere prefrontal TMS trains did not modulate sensor-space left prefrontal cortical excitability, but did modulate right frontal TEPs. 12 sequential left prefrontal TMS trains were not sufficient to induce cumulative neural changes. Exploratory analyses suggest train-induced non-local cortical excitability changes in the later parts of the TEP, changes in phase coordination dynamics, and distributed changes using source space models. This work provides important groundwork for future directions and highlights the experimental complexity required to measure acute neural changes to TMS treatment.

## Acknowledgments

We would like to acknowledge the contributions of all of our research participants. We extend gratitude to the members of the Personalized Neurotherapeutics Laboratory for helpful feedback on the article and throughout the course of the study. We would like to acknowledge Shannon Proksch, PhD, Augustana University for her useful comments on the RQA.

## Author contributions

JMR, CCC, and CJK conceptualized and designed the study, with significant contributions from MS and JT. CCC programmed the experiment. MS and JT collected the data. JMR, CCC, and MS conducted the analyses. All authors interpreted the results. All authors contributed to the writing of the manuscript. All authors provided intellectual contributions to and approval of the final manuscript.

## Availability of Data and Materials

The datasets generated and/or analyzed during the current study are available in the National Institute of Mental Health Data Archive linked to the project Closing the loop: development of real-time, personalized brain stimulation #4223 [ID 2263] [https://nda.nih.gov/experiment.html?id=2263&collectionId=4223]

## Supplementary Information

The online version contains supplementary material.

**Correspondence** and requests for materials should be addressed to C.J.K.

## References

[1] Chail A, Saini RK, Bhat PS, Srivastava K, Chauhan V. Transcranial magnetic stimulation: A review of its evolution and current applications. Ind Psychiatry J 2018;27:172–80. 10.4103/ipj.ipj_88_18.

[2] Blumberger DM, Vila-Rodriguez F, Thorpe KE, Feffer K, Noda Y, Giacobbe P, et al. Effectiveness of theta burst versus high-frequency repetitive transcranial magnetic stimulation in patients with depression (THREE-D): a randomised non-inferiority trial. Lancet 2018;391:1683–92. 10.1016/S0140-6736(18)30295-2.

[3] Trevizol AP, Downar J, Vila-Rodriguez F, Thorpe KE, Daskalakis ZJ, Blumberger DM. Predictors of remission after repetitive transcranial magnetic stimulation for the treatment of major depressive disorder: An analysis from the randomised non-inferiority THREE-D trial. EClinicalMedicine 2020;22:100349. 10.1016/j.eclinm.2020.100349.

[4] Stefanou M-I, Baur D, Belardinelli P, Bergmann TO, Blum C, Gordon PC, et al. Brain State-dependent Brain Stimulation with Real-time Electroencephalography-Triggered Transcranial Magnetic Stimulation. JoVE 2019:59711. 10.3791/59711.

[5] Zrenner B, Zrenner C, Gordon PC, Belardinelli P, McDermott EJ, Soekadar SR, et al. Brain oscillation-synchronized stimulation of the left dorsolateral prefrontal cortex in depression using real-time EEG-triggered TMS. Brain Stimulation 2020;13:197–205. 10.1016/j.brs.2019.10.007.

[6] Zrenner C, Desideri D, Belardinelli P, Ziemann U. Real-time EEG-defined excitability states determine efficacy of TMS-induced plasticity in human motor cortex. Brain Stimul 2018;11:374–89. 10.1016/j.brs.2017.11.016.

[7] Eshel N, Keller CJ, Wu W, Jiang J, Mills-Finnerty C, Huemer J, et al. Global connectivity and local excitability changes underlie antidepressant effects of repetitive transcranial magnetic stimulation. Neuropsychopharmacol 2020;45:1018–25. 10.1038/s41386-020-0633-z.

[8] Ilmoniemi RJ, Virtanen J, Ruohonen J, Karhu J, Aronen HJ, Näätänen R, et al. Neuronal responses to magnetic stimulation reveal cortical reactivity and connectivity: NeuroReport 1997;8:3537–40. 10.1097/00001756-199711100-00024.

[9] Ilmoniemi RJ, Kičić D. Methodology for Combined TMS and EEG. Brain Topogr 2010;22:233–48. 10.1007/s10548-009-0123-4.

[10] Ozdemir RA, Tadayon E, Boucher P, Sun H, Momi D, Ganglberger W, et al. Cortical responses to noninvasive perturbations enable individual brain fingerprinting. Brain Stimulation 2021;14:391–403. 10.1016/j.brs.2021.02.005.

[11] Rogasch NC, Fitzgerald PB. Assessing cortical network properties using TMS-EEG. Hum Brain Mapp 2013;34:1652–69. 10.1002/hbm.22016.

[12] Esser SK, Huber R, Massimini M, Peterson MJ, Ferrarelli F, Tononi G. A direct demonstration of cortical LTP in humans: A combined TMS/EEG study. Brain Research Bulletin 2006;69:86–94. 10.1016/j.brainresbull.2005.11.003.

[13] Hamidi M, Slagter HA, Tononi G, Postle BR. Brain responses evoked by high-frequency repetitive transcranial magnetic stimulation: an event-related potential study. Brain Stimul 2010;3:2–14. 10.1016/j.brs.2009.04.001.

[14] Veniero D, Maioli C, Miniussi C. Potentiation of short-latency cortical responses by high-frequency repetitive transcranial magnetic stimulation. J Neurophysiol 2010;104:1578–88. 10.1152/jn.00172.2010.

[15] Tremblay S, Rogasch NC, Premoli I, Blumberger DM, Casarotto S, Chen R, et al. Clinical utility and prospective of TMS-EEG. Clin Neurophysiol 2019;130:802–44. 10.1016/j.clinph.2019.01.001.

[16] Kähkönen S, Komssi S, Wilenius J, Ilmoniemi RJ. Prefrontal transcranial magnetic stimulation produces intensity-dependent EEG responses in humans. NeuroImage 2005;24:955–60. 10.1016/j.neuroimage.2004.09.048.

[17] Lioumis P, Kičić D, Savolainen P, Mäkelä JP, Kähkönen S. Reproducibility of TMS-Evoked EEG responses. Hum Brain Mapp 2009;30:1387–96. 10.1002/hbm.20608.

[18] Lucas MV, Cline CC, Sun Y, Yan M, Hogoboom N, Etkin A. Characterization of rTMS acute response profiles for systematic design of neuromodulation interventions. In revision.

[19] Koenigs M, Grafman J. The functional neuroanatomy of depression: distinct roles for ventromedial and dorsolateral prefrontal cortex. Behav Brain Res 2009;201:239–43. 10.1016/j.bbr.2009.03.004.

[20] Voineskos D, Blumberger DM, Rogasch NC, Zomorrodi R, Farzan F, Foussias G, et al. Neurophysiological effects of repetitive transcranial magnetic stimulation (rTMS) in treatment resistant depression. Clin Neurophysiol 2021;132:2306–16. 10.1016/j.clinph.2021.05.008.

[21] Voineskos D, Blumberger DM, Zomorrodi R, Rogasch NC, Farzan F, Foussias G, et al. Altered Transcranial Magnetic Stimulation-Electroencephalographic Markers of Inhibition and Excitation in the Dorsolateral Prefrontal Cortex in Major Depressive Disorder. Biol Psychiatry 2019;85:477–86. 10.1016/j.biopsych.2018.09.032.

[22] Keller CJ, Huang Y, Herrero JL, Fini ME, Du V, Lado FA, et al. Induction and Quantification of Excitability Changes in Human Cortical Networks. J Neurosci 2018;38:5384–98. 10.1523/JNEUROSCI.1088-17.2018.

[23] Biabani M, Fornito A, Mutanen TP, Morrow J, Rogasch NC. Characterizing and minimizing the contribution of sensory inputs to TMS-evoked potentials. Brain Stimul 2019;12:1537–52. 10.1016/j.brs.2019.07.009.

[24] Freedberg M, Reeves JA, Hussain SJ, Zaghloul KA, Wassermann EM. Identifying site-and stimulation-specific TMS-evoked EEG potentials using a quantitative cosine similarity metric. PLoS ONE 2020;15:e0216185. 10.1371/journal.pone.0216185.

[25] Rocchi L, Di Santo A, Brown K, Ibáñez J, Casula E, Rawji V, et al. Disentangling EEG responses to TMS due to cortical and peripheral activations. Brain Stimul 2021;14:4–18. 10.1016/j.brs.2020.10.011.

[26] Ross JM, Ozdemir RA, Lian SJ, Fried PJ, Schmitt EM, Inouye SK, et al. A structured ICA-based process for removing auditory evoked potentials. Sci Rep 2022;12:1391. 10.1038/s41598-022-05397-3.

[27] Ross JM, Sarkar M, Keller CJ. Experimental suppression of transcranial magnetic stimulation-electroencephalography sensory potentials. Hum Brain Mapp 2022;43:5141–53. 10.1002/hbm.25990.

[28] Richardson M, Paxton, Alexandra, Kuznetsov, Nikita. Nonlinear methods for understanding complex dynamical phenomena in psychological science. APA Psychological Science Agenda, 2017.

[29] Richardson MJ, Schmidt RC, Kay BA. Distinguishing the noise and attractor strength of coordinated limb movements using recurrence analysis. Biol Cybern 2007;96:59–78. 10.1007/s00422-006-0104-6.

[30] Marwan N. A historical review of recurrence plots. Eur Phys J Spec Top 2008;164:3–12. 10.1140/epjst/e2008-00829-1.

[31] Rossi S, Hallett M, Rossini PM, Pascual-Leone A. Screening questionnaire before TMS: An update. Clinical Neurophysiology 2011;122:1686. 10.1016/j.clinph.2010.12.037.

[32] Yeung A, Feldman G, Pedrelli P, Hails K, Fava M, Reyes T, et al. The Quick Inventory of Depressive Symptomatology, clinician rated and self-report: a psychometric assessment in Chinese Americans with major depressive disorder. J Nerv Ment Dis 2012;200:712–5. 10.1097/NMD.0b013e318261413d.

[33] Rush AJ, Trivedi MH, Ibrahim HM, Carmody TJ, Arnow B, Klein DN, et al. The 16-Item Quick Inventory of Depressive Symptomatology (QIDS), clinician rating (QIDS-C), and self-report (QIDS-SR): a psychometric evaluation in patients with chronic major depression. Biol Psychiatry 2003;54:573–83. 10.1016/s0006-3223(02)01866-8.

[34] Ilmoniemi RJ, Kičić D. Methodology for Combined TMS and EEG. Brain Topogr 2010;22:233–48. 10.1007/s10548-009-0123-4.

[35] Conde V, Tomasevic L, Akopian I, Stanek K, Saturnino GB, Thielscher A, et al. The non-transcranial TMS-evoked potential is an inherent source of ambiguity in TMS-EEG studies. NeuroImage 2019;185:300–12. 10.1016/j.neuroimage.2018.10.052.

[36] Nikouline V, Ruohonen J, Ilmoniemi RJ. The role of the coil click in TMS assessed with simultaneous EEG. Clinical Neurophysiology 1999;110:1325–8. 10.1016/S1388-2457(99)00070-X.

[37] Gordon PC, Desideri D, Belardinelli P, Zrenner C, Ziemann U. Comparison of cortical EEG responses to realistic sham versus real TMS of human motor cortex. Brain Stimulation 2018;11:1322–30. 10.1016/j.brs.2018.08.003.

[38] Rossi S, Hallett M, Rossini PM, Pascual-Leone A, Safety of TMS Consensus Group. Safety, ethical considerations, and application guidelines for the use of transcranial magnetic stimulation in clinical practice and research. Clin Neurophysiol 2009;120:2008–39. 10.1016/j.clinph.2009.08.016.

[39] Rossini PM, Barker AT, Berardelli A, Caramia MD, Caruso G, Cracco RQ, et al. Non-invasive electrical and magnetic stimulation of the brain, spinal cord and roots: basic principles and procedures for routine clinical application. Report of an IFCN committee. Electroencephalography and Clinical Neurophysiology 1994;91:79–92. 10.1016/0013-4694(94)90029-9.

[40] Rossini PM, Burke D, Chen R, Cohen LG, Daskalakis Z, Di Iorio R, et al. Non-invasive electrical and magnetic stimulation of the brain, spinal cord, roots and peripheral nerves: Basic principles and procedures for routine clinical and research application. An updated report from an I.F.C.N. Committee. Clin Neurophysiol 2015;126:1071–107. 10.1016/j.clinph.2015.02.001.

[41] Stokes MG, Chambers CD, Gould IC, Henderson TR, Janko NE, Allen NB, et al. Simple Metric For Scaling Motor Threshold Based on Scalp-Cortex Distance: Application to Studies Using Transcranial Magnetic Stimulation. Journal of Neurophysiology 2005;94:4520–7. 10.1152/jn.00067.2005.

[42] Pridmore S, Fernandes Filho JA, Nahas Z, Liberatos C, George MS. Motor threshold in transcranial magnetic stimulation: a comparison of a neurophysiological method and a visualization of movement method. J ECT 1998;14:25–7.

[43] Chen AC, Oathes DJ, Chang C, Bradley T, Zhou Z-W, Williams LM, et al. Causal interactions between fronto-parietal central executive and default-mode networks in humans. Proceedings of the National Academy of Sciences 2013;110:19944–9. 10.1073/pnas.1311772110.

[44] Nielsen JD, Madsen KH, Puonti O, Siebner HR, Bauer C, Madsen CG, et al. Automatic skull segmentation from MR images for realistic volume conductor models of the head: Assessment of the state-of-the-art. Neuroimage 2018;174:587–98. 10.1016/j.neuroimage.2018.03.001.

[45] Thielscher A, Antunes A, Saturnino GB. Field modeling for transcranial magnetic stimulation: A useful tool to understand the physiological effects of TMS? 2015 37th Annual International Conference of the IEEE Engineering in Medicine and Biology Society (EMBC), Milan: IEEE; 2015, p. 222–5. 10.1109/EMBC.2015.7318340.

[46] Li X, Hartwell KJ, Owens M, Lematty T, Borckardt JJ, Hanlon CA, et al. Repetitive transcranial magnetic stimulation of the dorsolateral prefrontal cortex reduces nicotine cue craving. Biol Psychiatry 2013;73:714–20. 10.1016/j.biopsych.2013.01.003.

[47] Li X, Du L, Sahlem GL, Badran BW, Henderson S, George MS. Repetitive transcranial magnetic stimulation (rTMS) of the dorsolateral prefrontal cortex reduces resting-state insula activity and modulates functional connectivity of the orbitofrontal cortex in cigarette smokers. Drug Alcohol Depend 2017;174:98–105. 10.1016/j.drugalcdep.2017.02.002.

[48] Liu Q, Sun H, Hu Y, Wang Q, Zhao Z, Dong D, et al. Intermittent Theta Burst Stimulation vs. High-Frequency Repetitive Transcranial Magnetic Stimulation in the Treatment of Methamphetamine Patients. Front Psychiatry 2022;13:842947. 10.3389/fpsyt.2022.842947.

[49] Shen Y, Cao X, Tan T, Shan C, Wang Y, Pan J, et al. 10-Hz Repetitive Transcranial Magnetic Stimulation of the Left Dorsolateral Prefrontal Cortex Reduces Heroin Cue Craving in Long-Term Addicts. Biol Psychiatry 2016;80:e13–14. 10.1016/j.biopsych.2016.02.006.

[50] Gordon PC, Jovellar DB, Song Y, Zrenner C, Belardinelli P, Siebner HR, et al. Recording brain responses to TMS of primary motor cortex by EEG - utility of an optimized sham procedure. Neuroimage 2021;245:118708. 10.1016/j.neuroimage.2021.118708.

[51] Veniero D, Bortoletto M, Miniussi C. TMS-EEG co-registration: On TMS-induced artifact. Clinical Neurophysiology 2009;120:1392–9. 10.1016/j.clinph.2009.04.023.

[52] Delorme A, Makeig S. EEGLAB: an open source toolbox for analysis of single-trial EEG dynamics including independent component analysis. Journal of Neuroscience Methods 2004;134:9–21. 10.1016/j.jneumeth.2003.10.009.

[53] Cline CC, Lucas MV, Sun Y, Menezes M, Etkin A. Advanced Artifact Removal for Automated TMS-EEG Data Processing. 2021 10th International IEEE/EMBS Conference on Neural Engineering (NER), Italy: IEEE; 2021, p. 1039–42. 10.1109/NER49283.2021.9441147.

[54] Rogasch NC, Biabani M, Mutanen TP. Designing and comparing cleaning pipelines for TMS-EEG data: A theoretical overview and practical example. Journal of Neuroscience Methods 2022;371:109494. 10.1016/j.jneumeth.2022.109494.

[55] Pion-Tonachini L, Kreutz-Delgado K, Makeig S. ICLabel: An automated electroencephalographic independent component classifier, dataset, and website. Neuroimage 2019;198:181–97. 10.1016/j.neuroimage.2019.05.026.

[56] Mutanen TP, Metsomaa J, Liljander S, Ilmoniemi RJ. Automatic and robust noise suppression in EEG and MEG: The SOUND algorithm. NeuroImage 2018;166:135–51. 10.1016/j.neuroimage.2017.10.021.

[57] Beauchemin M, De Beaumont L. Statistical analysis of the mismatch negativity: To a dilemma, an answer. TQMP 2005;1:18–24. 10.20982/tqmp.01.1.p018.

[58] Kappenman ES, Luck SJ. Best Practices for Event-Related Potential Research in Clinical Populations. Biol Psychiatry Cogn Neurosci Neuroimaging 2016;1:110–5. 10.1016/j.bpsc.2015.11.007.

[59] Gogulski J, Cline CC, Ross JM, Truong J, Sarkar M, Parmigiani S, et al. Mapping cortical excitability in the human dorsolateral prefrontal cortex. bioRxiv 2023. 10.1101/2023.01.20.524867.

[60] Pernet CR, Chauveau N, Gaspar C, Rousselet GA. LIMO EEG: a toolbox for hierarchical LInear MOdeling of ElectroEncephaloGraphic data. Comput Intell Neurosci 2011;2011:831409. 10.1155/2011/831409.

[61] Proksch S, Reeves M, Spivey M, Balasubramaniam R. Coordination dynamics of multi-agent interaction in a musical ensemble. Sci Rep 2022;12:421. 10.1038/s41598-021-04463-6.

[62] Marwan N, Webber CL. Mathematical and Computational Foundations of Recurrence Quantifications. In: Webber, CL, Marwan N, editors. Recurrence Quantification Analysis, Cham: Springer International Publishing; 2015, p. 3–43. 10.1007/978-3-319-07155-8_1.

[63] Scheurich R, Demos AP, Zamm A, Mathias B, Palmer C. Capturing Intra-and Inter-Brain Dynamics with Recurrence Quantification Analysis. 41st Annual Meeting of the Cognitive Science Society 2019:2748–54.

[64] Ross JM, Balasubramaniam R. Auditory white noise reduces postural fluctuations even in the absence of vision. Exp Brain Res 2015;233:2357–63. 10.1007/s00221-015-4304-y.

[65] Abney DH, Warlaumont AS, Haussman A, Ross JM, Wallot S. Using nonlinear methods to quantify changes in infant limb movements and vocalizations. Front Psychol 2014;5:771. 10.3389/fpsyg.2014.00771.

[66] Kelso JAS. Multistability and metastability: understanding dynamic coordination in the brain. Philos Trans R Soc Lond B Biol Sci 2012;367:906–18. 10.1098/rstb.2011.0351.

[67] Michel CM, Murray MM. Towards the utilization of EEG as a brain imaging tool. NeuroImage 2012;61:371–85. 10.1016/j.neuroimage.2011.12.039.

[68] Gramfort A, Papadopoulo T, Olivi E, Clerc M. OpenMEEG: opensource software for quasistatic bioelectromagnetics. BioMed Eng OnLine 2010;9:45. 10.1186/1475-925X-9-45.

[69] Kybic J, Clerc M, Abboud T, Faugeras O, Keriven R, Papadopoulo T. A common formalism for the Integral formulations of the forward EEG problem. IEEE Trans Med Imaging 2005;24:12–28. 10.1109/TMI.2004.837363.

[70] Tadel F, Baillet S, Mosher JC, Pantazis D, Leahy RM. Brainstorm: A User-Friendly Application for MEG/EEG Analysis. Computational Intelligence and Neuroscience 2011;2011:1–13. 10.1155/2011/879716.

[71] Pistorius T, Aldrich C, Auret L, Pineda J. Early Detection of risk of autism spectrum disorder based on recurrence quantification analysis of electroencephalographic signals. 2013 6th International IEEE/EMBS Conference on Neural Engineering (NER), San Diego, CA, USA: IEEE; 2013, p. 198–201. 10.1109/NER.2013.6695906.

[72] Becker K, Schneider G, Eder M, Ranft A, Kochs EF, Zieglgänsberger W, et al. Anaesthesia Monitoring by Recurrence Quantification Analysis of EEG Data. PLoS ONE 2010;5:e8876. 10.1371/journal.pone.0008876.

[73] Hadley D, Anderson BS, Borckardt JJ, Arana A, Li X, Nahas Z, et al. Safety, tolerability, and effectiveness of high doses of adjunctive daily left prefrontal repetitive transcranial magnetic stimulation for treatment-resistant depression in a clinical setting. J ECT 2011;27:18–25. 10.1097/YCT.0b013e3181ce1a8c.

[74] Avery DH, Isenberg KE, Sampson SM, Janicak PG, Lisanby SH, Maixner DF, et al. Transcranial magnetic stimulation in the acute treatment of major depressive disorder: clinical response in an open-label extension trial. J Clin Psychiatry 2008;69:441–51. 10.4088/jcp.v69n0315.

[75] Yip AG, George MS, Tendler A, Roth Y, Zangen A, Carpenter LL. 61% of unmedicated treatment resistant depression patients who did not respond to acute TMS treatment responded after four weeks of twice weekly deep TMS in the Brainsway pivotal trial. Brain Stimul 2017;10:847–9. 10.1016/j.brs.2017.02.013.

[76] Jung SH, Shin JE, Jeong Y-S, Shin H-I. Changes in motor cortical excitability induced by high-frequency repetitive transcranial magnetic stimulation of different stimulation durations. Clinical Neurophysiology 2008;119:71–9. 10.1016/j.clinph.2007.09.124.

[77] Jin J, Wang X, Wang H, Li Y, Liu Z, Yin T. Train duration and inter-train interval determine the direction and intensity of high-frequency rTMS after-effects. Front Neurosci 2023;17:1157080. 10.3389/fnins.2023.1157080.

[78] Quartarone A, Bagnato S, Rizzo V, Morgante F, Sant’Angelo A, Battaglia F, et al. Distinct changes in cortical and spinal excitability following high-frequency repetitive TMS to the human motor cortex. Exp Brain Res 2005;161:114–24. 10.1007/s00221-004-2052-5.

[79] Peinemann A, Reimer B, Löer C, Quartarone A, Münchau A, Conrad B, et al. Long-lasting increase in corticospinal excitability after 1800 pulses of subthreshold 5 Hz repetitive TMS to the primary motor cortex. Clinical Neurophysiology 2004;115:1519–26. 10.1016/j.clinph.2004.02.005.

[80] Ye Y, Wang J, Che X. Concurrent TMS-EEG to reveal the neuroplastic changes in the prefrontal and insular cortices in the analgesic effects of DLPFC-rTMS. Cereb Cortex 2022;32:4436–46. 10.1093/cercor/bhab493.

[81] Du X, Rowland LM, Summerfelt A, Choa F-S, Wittenberg GF, Wisner K, et al. Cerebellar-Stimulation Evoked Prefrontal Electrical Synchrony Is Modulated by GABA. Cerebellum 2018;17:550–63. 10.1007/s12311-018-0945-2.

[82] Ferrarelli F, Massimini M, Sarasso S, Casali A, Riedner BA, Angelini G, et al. Breakdown in cortical effective connectivity during midazolam-induced loss of consciousness. Proceedings of the National Academy of Sciences 2010;107:2681–6. 10.1073/pnas.0913008107.

[83] Premoli I, Castellanos N, Rivolta D, Belardinelli P, Bajo R, Zipser C, et al. TMS-EEG signatures of GABAergic neurotransmission in the human cortex. J Neurosci 2014;34:5603–12. 10.1523/JNEUROSCI.5089-13.2014.

[84] Heunis T, Aldrich C, Peters JM, Jeste SS, Sahin M, Scheffer C, et al. Recurrence quantification analysis of resting state EEG signals in autism spectrum disorder – a systematic methodological exploration of technical and demographic confounders in the search for biomarkers. BMC Med 2018;16:101. 10.1186/s12916-018-1086-7.

[85] Bhat S, Acharya UR, Adeli H, Bairy GM, Adeli A. Automated diagnosis of autism: in search of a mathematical marker. Reviews in the Neurosciences 2014;25. 10.1515/revneuro-2014-0036.

[86] Acharya UR, Sree SV, Chattopadhyay S, Yu W, Ang PCA. Application of recurrence quantification analysis for the automated identification of epileptic EEG signals. Int J Neur Syst 2011;21:199–211. 10.1142/S0129065711002808.

[87] Song I-H, Lee D-S, Kim SI. Recurrence quantification analysis of sleep electoencephalogram in sleep apnea syndrome in humans. Neuroscience Letters 2004;366:148–53. 10.1016/j.neulet.2004.05.025.

[88] Siebner HR, Funke K, Aberra AS, Antal A, Bestmann S, Chen R, et al. Transcranial magnetic stimulation of the brain: What is stimulated? – A consensus and critical position paper. Clinical Neurophysiology 2022;140:59–97. 10.1016/j.clinph.2022.04.022.

[89] Parmigiani S, Ross JM, Cline C, Minasi C, Gogulski J, Keller CJ. Reliability and validity of TMS-EEG biomarkers. Biol Psychiatry: Cogn Neurosci 2022:S2451902222003408. 10.1016/j.bpsc.2022.12.005.

