## Supplement for "Neural effects of TMS trains on the human prefrontal cortex"

**Table S1 | Demographics.** (N=16)

| | |
|---|---|
| Age, mean years (SD) | 43.1 (12.5) |
| Sex | |
| Female, n (%) | 8 (50.0) |
| Male, n (%) | 8 (50.0) |
| Other or prefer not to state, n (%) | 0 (0.0) |
| Handedness | |
| Left hand dominant, n (%) | 1 (6.3) |
| Right hand dominant, n (%) | 15 (93.7) |
| Ambidextrous, n (%) | 0 (0.0) |
| Education | |
| GED or High School Diploma, n (%) | 0 (0.0) |
| Some college, no degree, n (%) | 0 (0.0) |
| Two year degree, n (%) | 2 (12.5) |
| Four year degree, n (%) | 8 (50.0) |
| Post graduate degree, n (%) | 6 (37.5) |
| Employment | |
| Part-time, n (%) | 3 (18.7) |
| Full-time, n (%) | 7 (43.7) |
| Unemployed, n (%) | 5 (31.3) |
| Retired, n (%) | 0 (0.0) |
| Full-time student, n (%) | 1 (6.3) |
| Race | |
| White, n (%) | 7 (43.7) |
| Black or African American, n (%) | 1 (6.3) |
| American Indian or Alaska Native, n (%) | 1 (6.3) |
| Asian, n (%) | 5 (31.2) |
| Native Hawaiian or Other Pacific Islander, n (%) | 0 (0.0) |
| Two or more races, n (%) | 0 (0.0) |
| Some other race or prefer not to state, n (%) | 2 (12.5) |

**Table S2 | Optimized coil angle and TMS train intensity for each subject.**

| | Angle (° from midline) | Intensity (% RMT) |
|---|---|---|
| S1 | 45 | 110 |
| S2 | 45 | 100 |
| S3 | 0 | 110 |
| S4 | 45 | 70 |
| S5 | 90 | 110 |
| S6 | 45 | 110 |
| S7 | 45 | 80 |
| S8 | 0 | 75 |
| S9 | 45 | 110 |
| S10 | 45 | 80 |
| S11 | 90 | 105 |
| S12 | 45 | 85 |
| S13 | 45 | 110 |
| S14 | 60 | 110 |
| S15 | 45 | 100 |
| S16 | 0 | 75 |

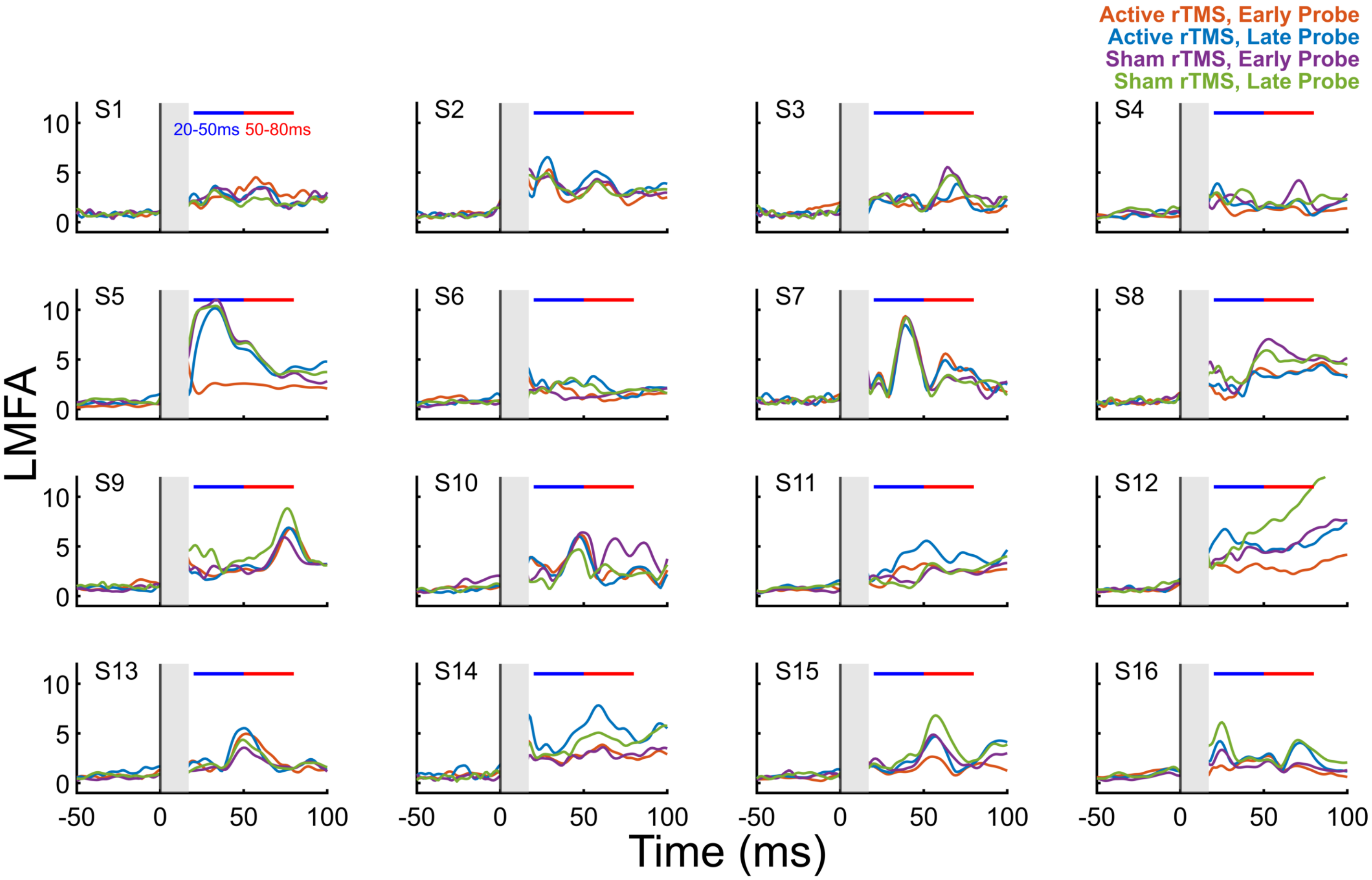

**Figure S1 | Individual subject LMFA time series in the four conditions.** The two analysis latency windows are indicated with a blue (20-50 ms) and a red (50-80 ms) horizontal bar. Time = 0 is the time of the single TMS probe pulse. For group average, see Fig 1D. For up to 300 ms, see Fig S2.

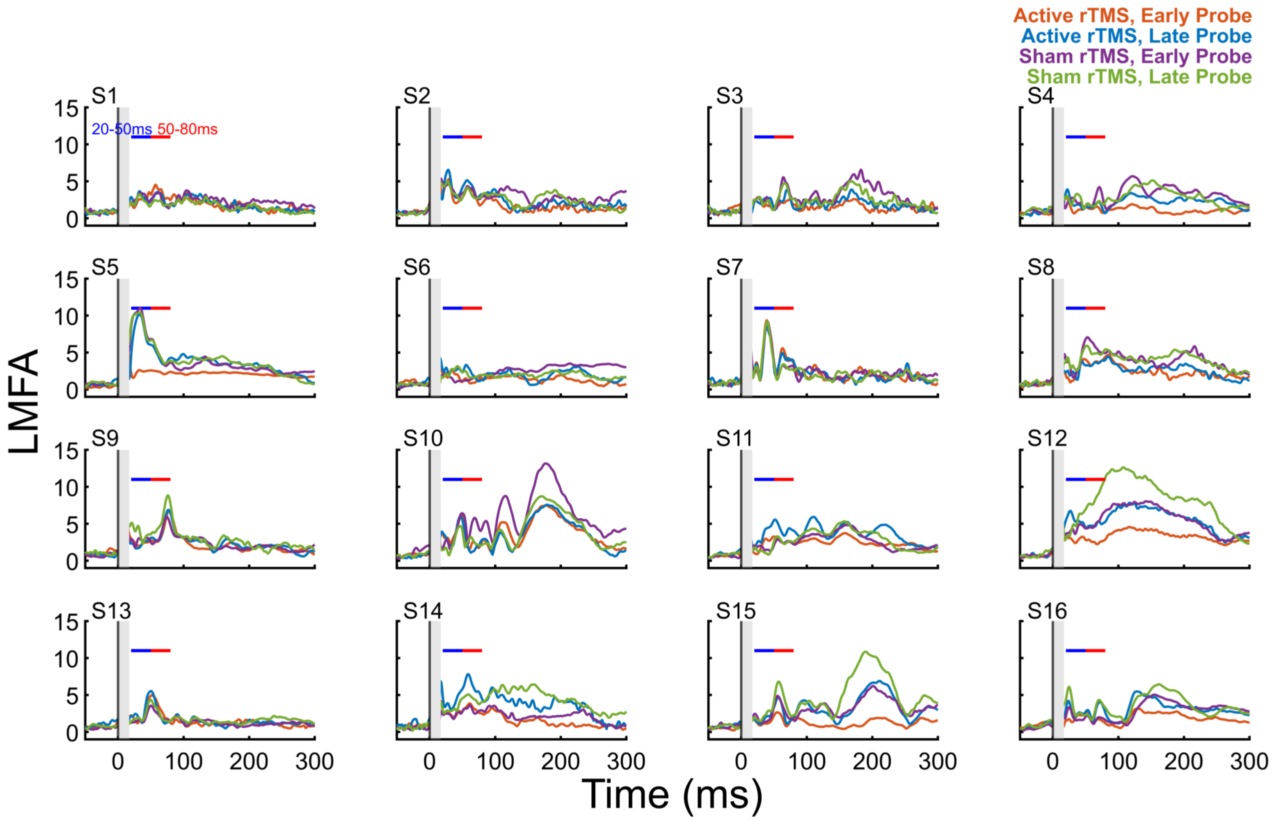

**Figure S2 | Individual subject LMFA time series from -50 to 300 ms.** For group average, see Fig 1D. For early TEP focusing on analysis latency windows, see Fig S1.

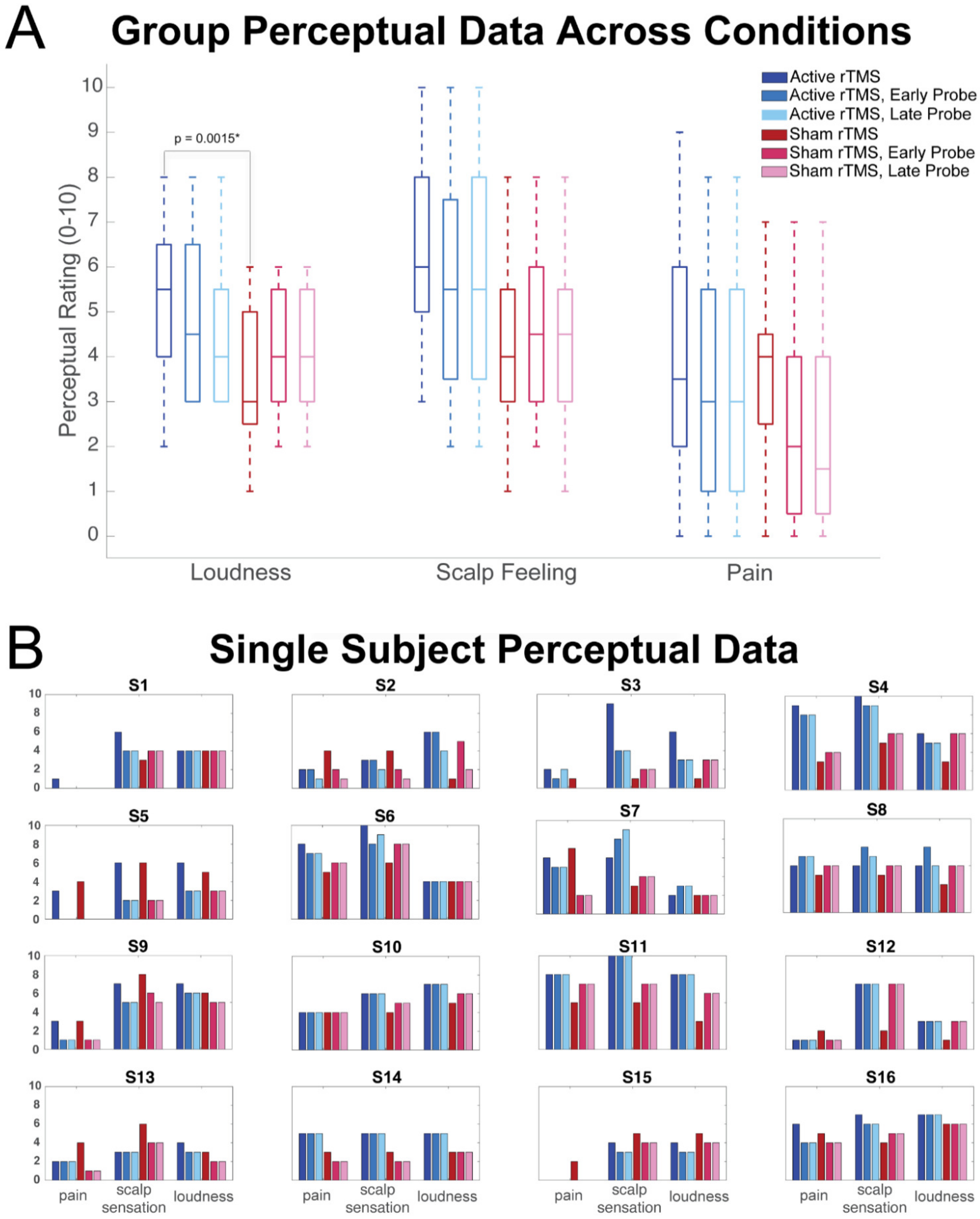

**Figure S3 | Perceptual ratings of sound loudness, scalp feeling, and pain.** A) Raw subject perceptual ratings for different stimulation conditions and latencies. B) Group-level perceptual rating averages for sham and active trains and probe latencies. Pairwise comparisons revealed that TMS sound loudness perception differed between active and sham trains ($p = 0.0015$, indicated with a bracket). We find no other significant differences between sham and active perceptual ratings.

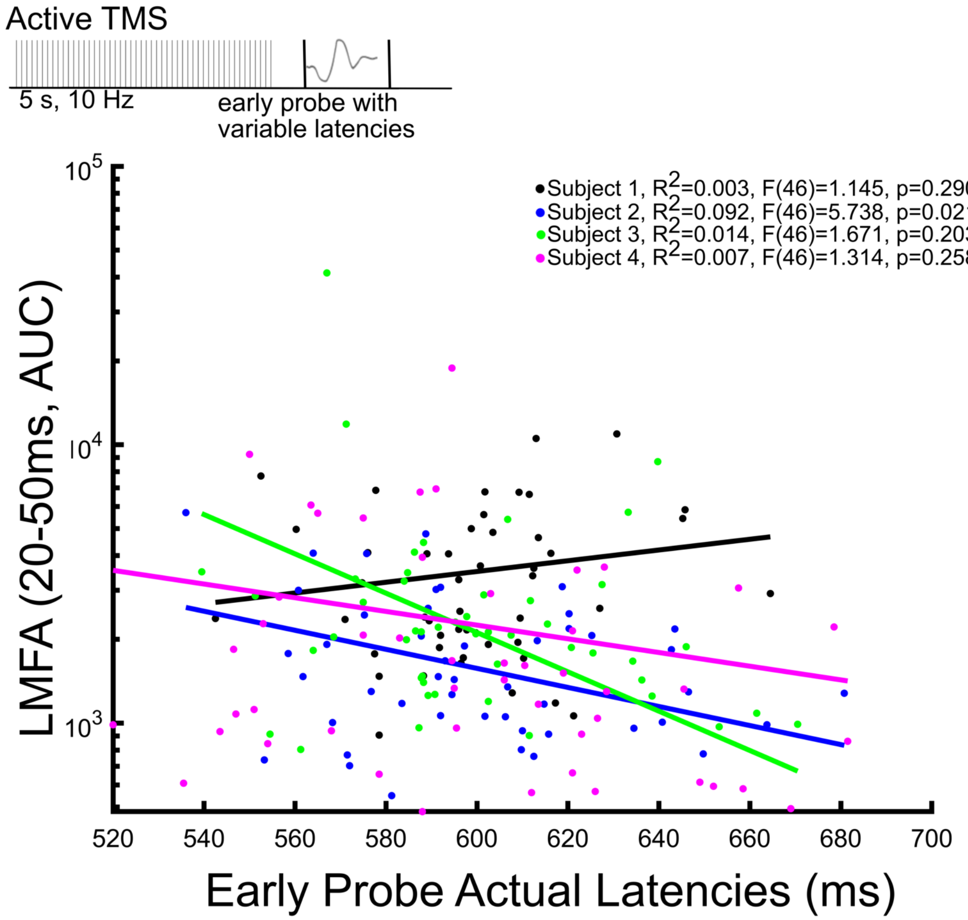

**Figure S4 | Exact timing of early probe latencies and LMFA (N=4).** Four subjects with early probe latencies ranging from 500-700 ms were examined in order to explore whether a linear relationship existed between exact early probe latency and LMFA AUC. Three of the four subjects trend toward a reduction in LMFA with later probe latencies, with one subject reaching significance. However, R-Squared values were very low indicating a weak linear relationship. No individual subject slopes were significantly different from 0 (subject 1: $t(47)=6.28$, $p=0.10$; subject 2: $t(47)=1.48$, $p=0.38$; subject 3: $t(47)=1.02$, $p=0.49$; subject 4: $t(47)=1.68$, $p=0.34$), further supporting that there does not appear to be a relationship between latency and LMFA.

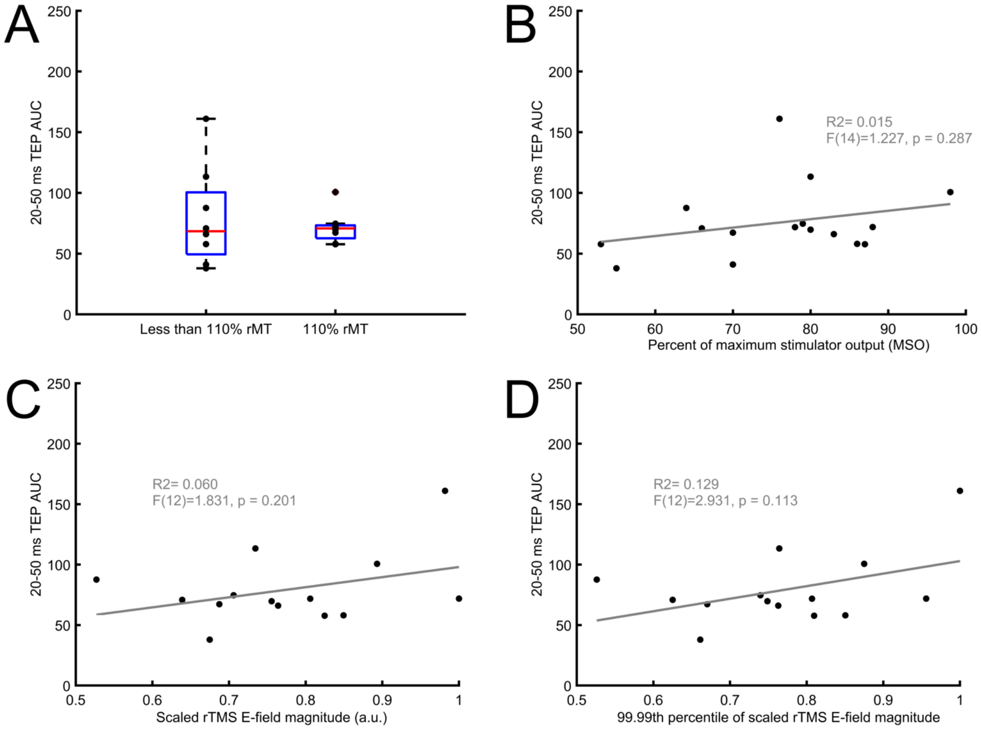

**Figure S5 | Relationship between TMS train intensity and early-local TEP.** A) The size of the early local TEP was comparable between subjects who received TMS trains at 110% RMT and subjects who needed TMS train intensity lowered (n=16). TEP size was not significantly predicted by B) % MSO (n=16), C) E-field (n=14), or D) the 99.99th percentile of the E-field (to avoid sensitivity to a very small number of single-vertex E-field peaks in the simulation results).

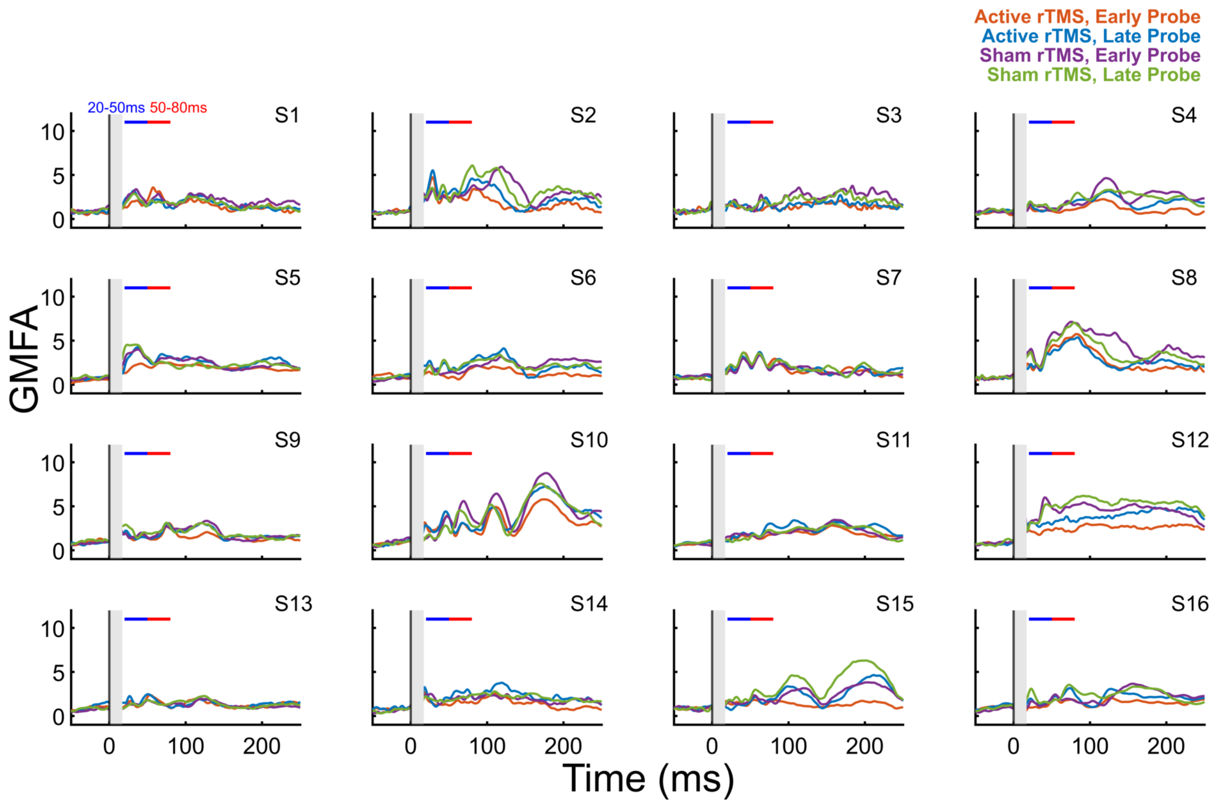

**Figure S6 | Individual subject GMFA time series in the four conditions.** All EEG channels are included in these averages. The two analysis latency windows are indicated with a blue (20-50 ms) and a red (50-80 ms) horizontal bar. Time = 0 is the time of the single TMS probe pulse.

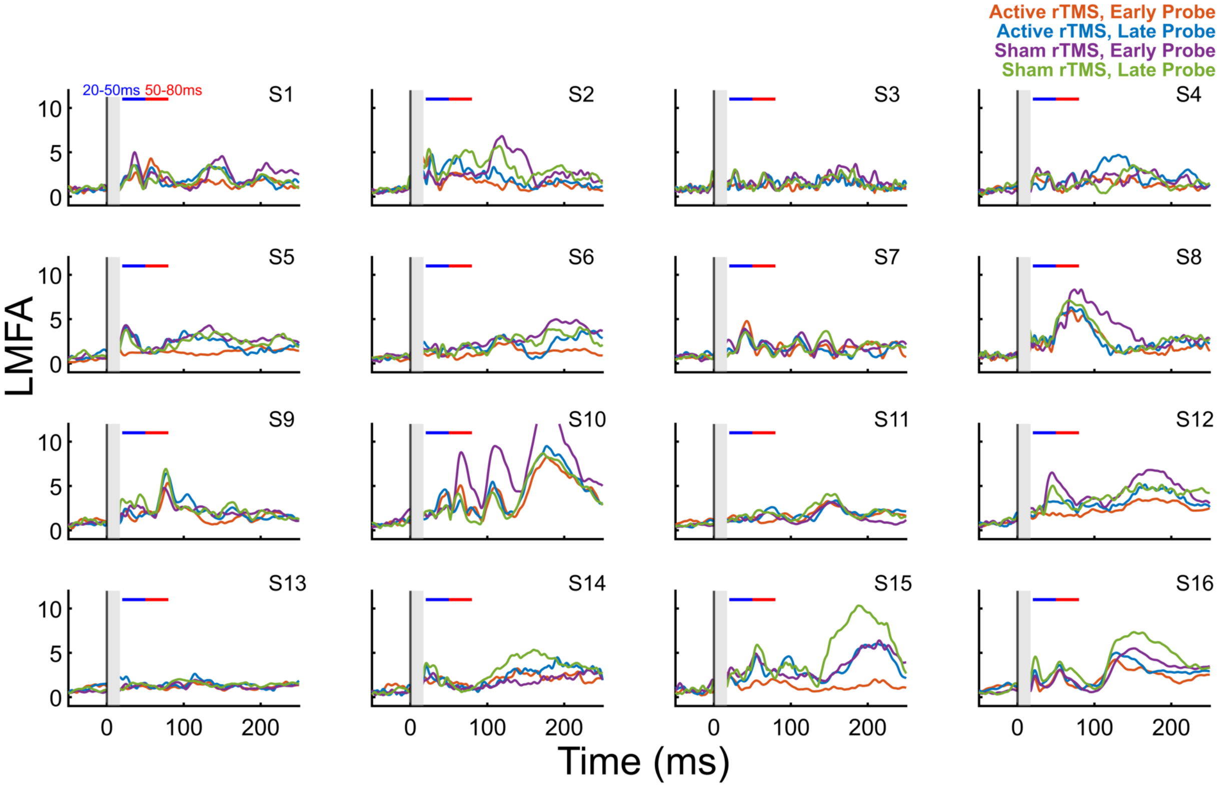

**Figure S7 | Individual subject right lateral LMFA time series in the four conditions.** The two analysis latency windows are indicated with a blue (20-50 ms) and a red (50-80 ms) horizontal bar. Time = 0 is the time of the single TMS probe pulse.

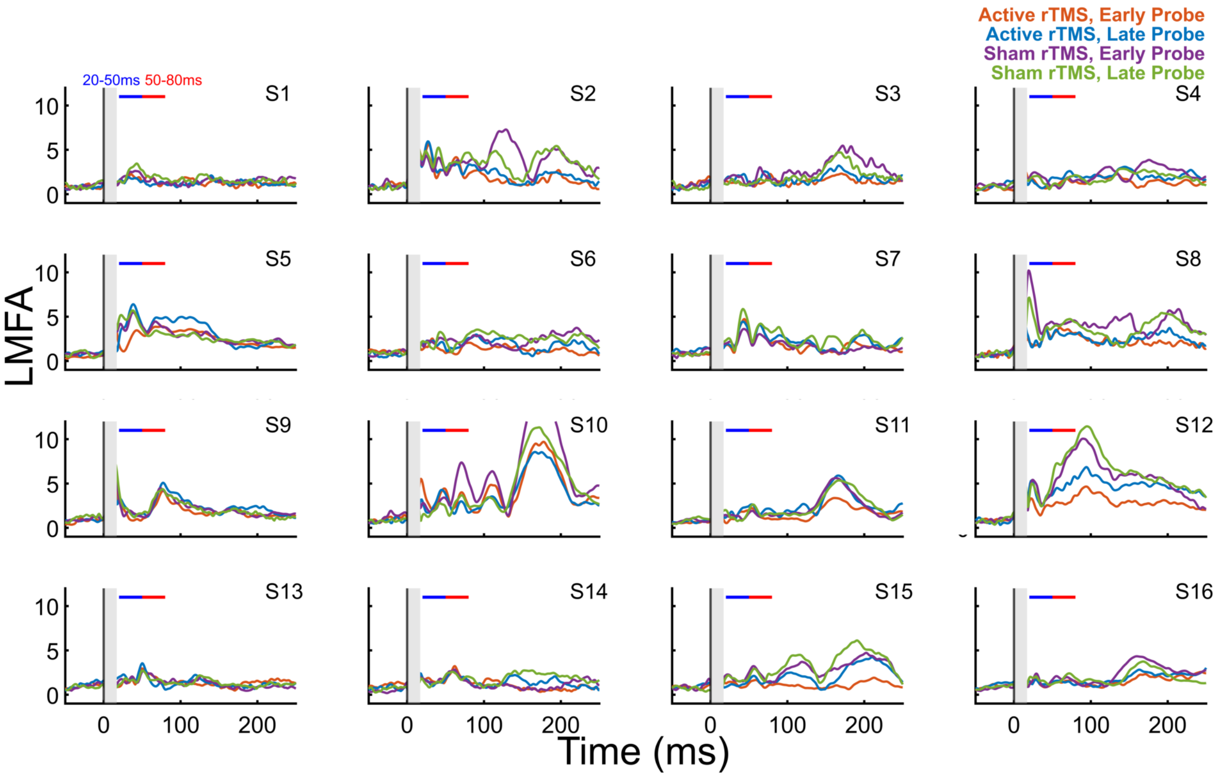

**Figure S8 | Individual subject left parietal LMFA time series in the four conditions.** The two analysis latency windows are indicated with a blue (20-50 ms) and a red (50-80 ms) horizontal bar. Time = 0 is the time of the single TMS probe pulse.

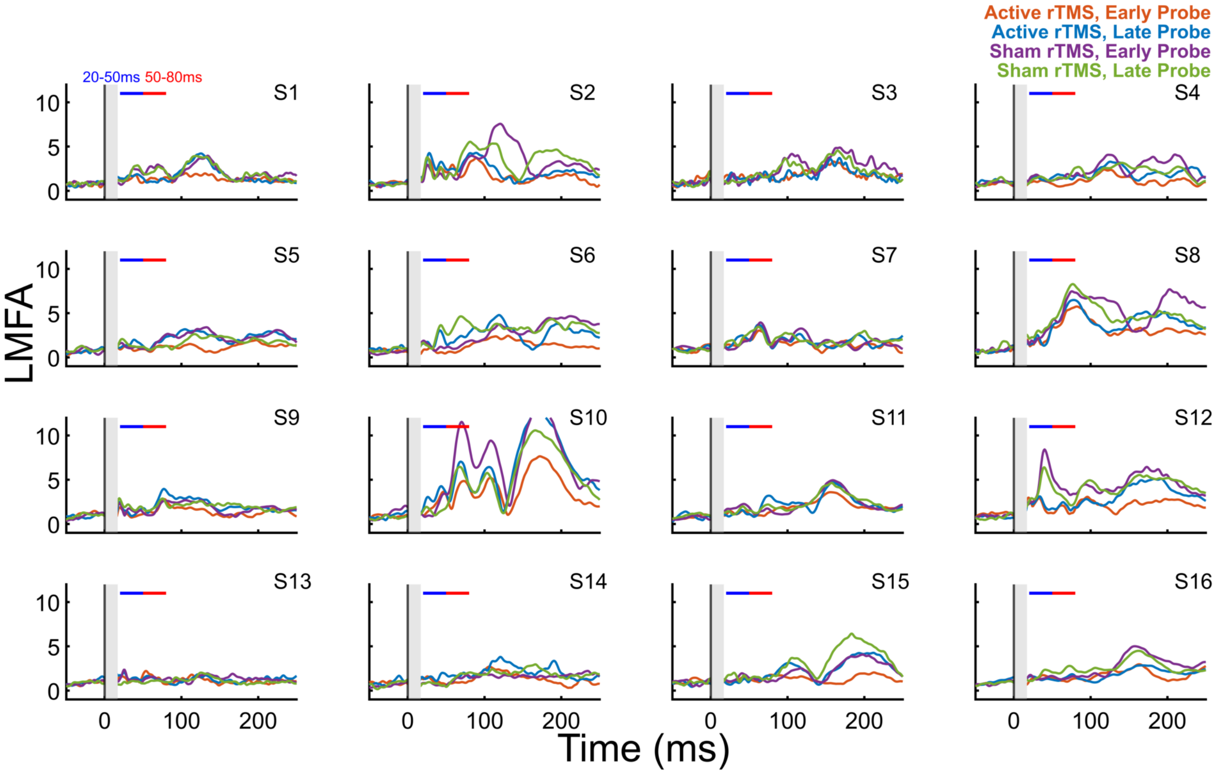

**Figure S9 | Individual subject right parietal LMFA time series in the four conditions.** The two analysis latency windows are indicated with a blue (20-50 ms) and a red (50-80 ms) horizontal bar. Time = 0 is the time of the single TMS probe pulse.

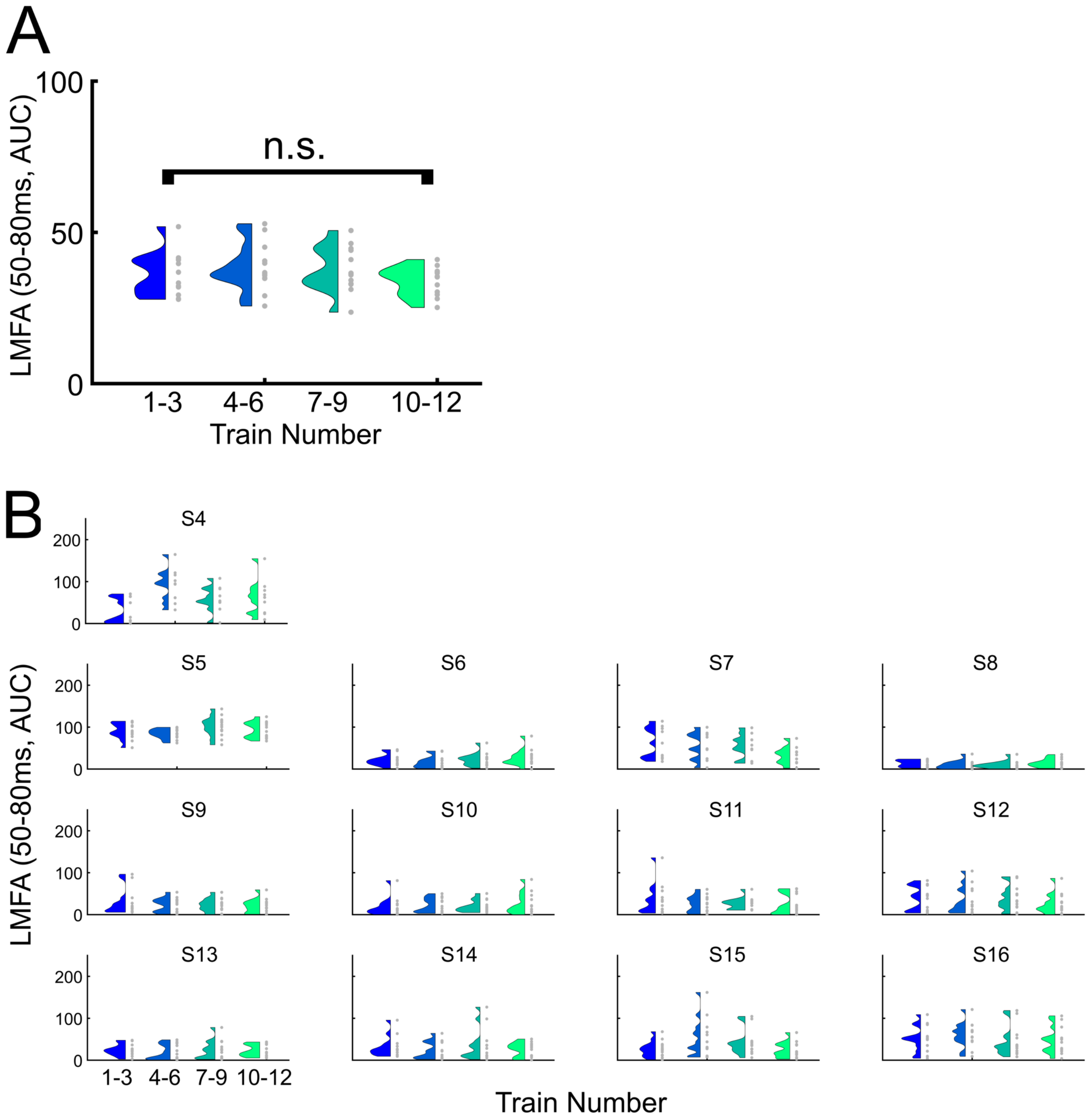

**Figure S10 | Sequential TMS trains did not modulate local TEPs at 50-80 ms latencies.** A) Group average (N=13) LMFA AUC from 50-80 ms latency window from the early probe following each sequential active TMS train, by groupings of 3 trains. Train averages across subjects in gray and distributions in color. There was no significant change in LMFA AUC across the four groups of trains, $F(3,36)=0.7722$, $p=0.5172$. B) Individual subject (averaged across 4 blocks) LMFA AUC from 50-80 ms latency window from the early probe following each sequential active TMS train in gray and distributions in color, by groupings of 3 trains.

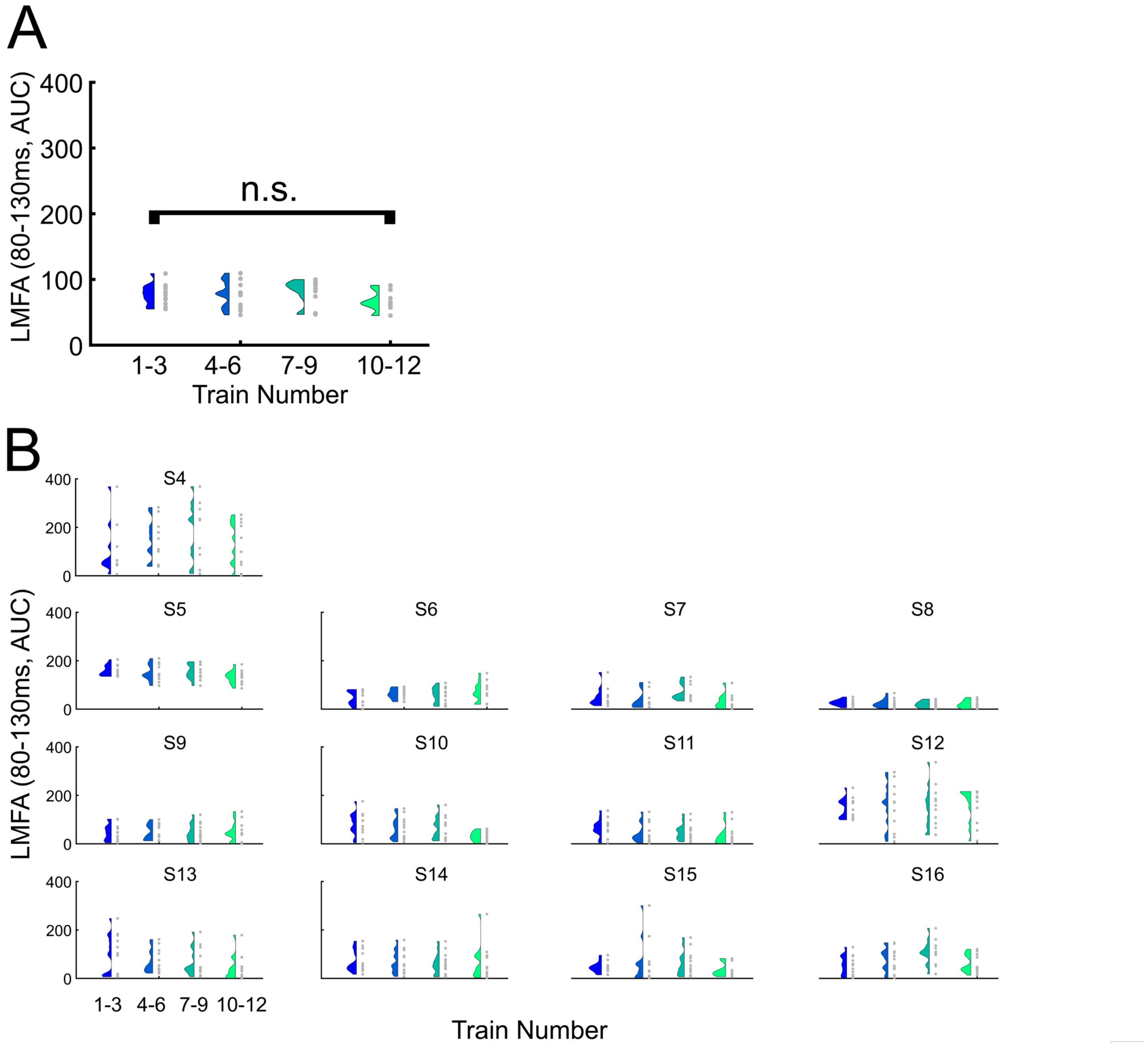

**Figure S11 | Sequential TMS trains did not modulate local TEPs at 80-130 ms latencies.** A) Group average (N=13) LMFA AUC from 80-130 ms latency window from the early probe following each sequential active TMS train, by groupings of 3 trains. Train averages across subjects in gray and distributions in color. There was no significant change in LMFA AUC across the four groups of trains, $F(3,36)=2.6313$, $p=0.0648$. B) Individual subject (averaged across 4 blocks) LMFA AUC from 80-130 ms latency window from the early probe following each sequential active TMS train in gray and distributions in color, by groupings of 3 trains.

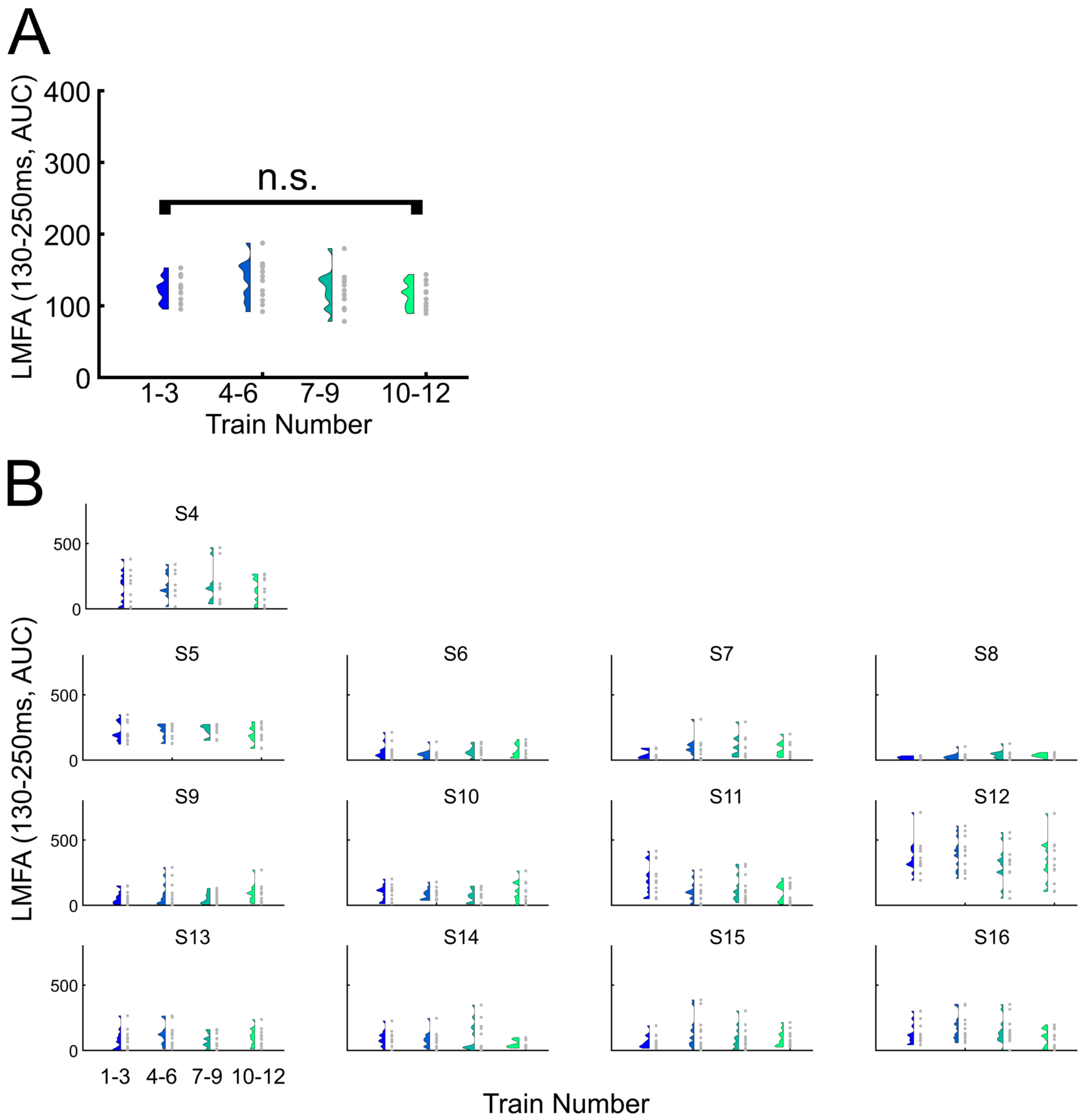

**Figure S12 | Sequential TMS trains did not modulate local TEPs at 130-250 ms latencies.** A) Group average (N=13) LMFA AUC from 130-250 ms latency window from the early probe following each sequential active TMS train, by groupings of 3 trains. Train averages across subjects in gray and distributions in color. There was no significant change in LMFA AUC across the four groups of trains, $F(3,36)=1.2487$, $p=0.3065$. B) Individual subject (averaged across 4 blocks) LMFA AUC from 130-250 ms latency window from the early probe following each sequential active TMS train in gray and distributions in color, by groupings of 3 trains.

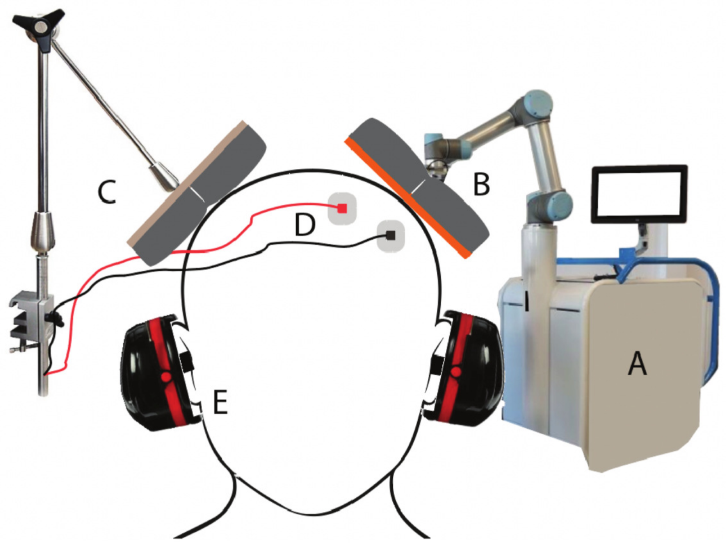

**Figure S13 | rTMS setup.** Active rTMS was administered using A) an Axilum Cobot holding a B) B65 coil with foam padding. Sham rTMS was administered using C) a flipped coil producing an auditory click from the TMS pulse as well as D) electrical stimulation to the left frontalis muscle. E) White noise auditory masking was played using earbud earplugs covered with over-the-ear noise minimizing headphones.

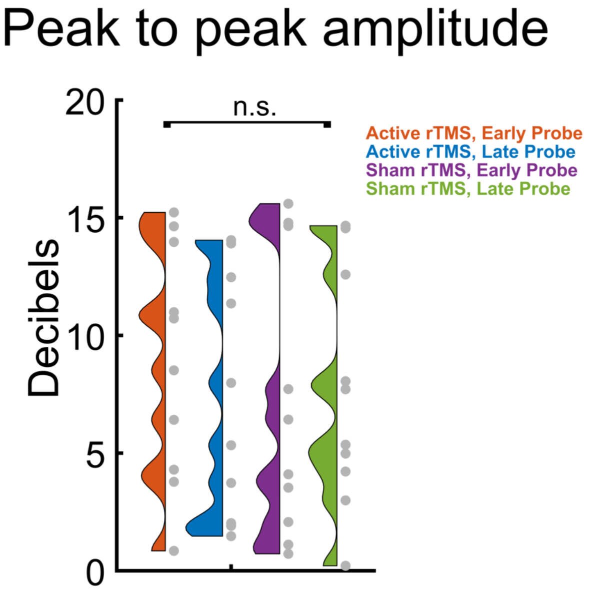

**Figure S14 | TMS trains did not modulate early local TEPs when using a peak to peak amplitude metric.** To verify that the results of the early local TEP analysis was not different when using peak to peak amplitudes rather than LMFA (Fig 1), we repeated the analysis using peak to peak amplitudes (in dBuV) for subjects with a negative peak between 20-50 ms and a positive peak between 50-80 ms. N=10 subjects fit these criteria. Shown are individual subject peak to peak amplitudes in the four conditions. We observed no main effect of probe latency ($F(1,9)=0.5255$, $p=0.4869$) and no main effect of stimulation ($F(1,9)=0.9619$, $p=0.3523$). Statistical test: 2 probe latencies (early, late) x 2 conditions (active, sham rTMS) repeated measures ANOVA.

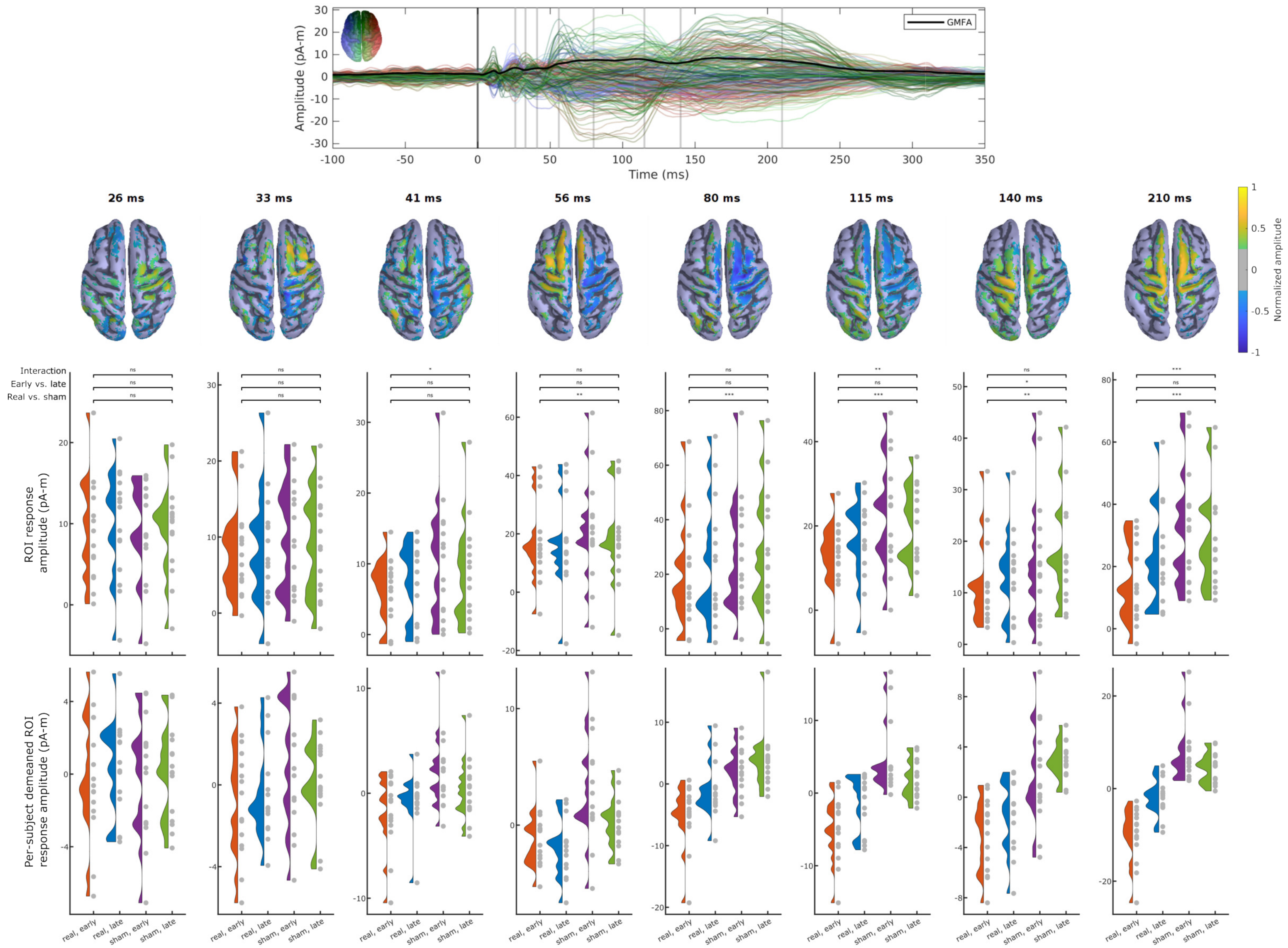

**Figure S15 | Source estimates across both early and late TEPs.** Source estimates for condition specific TEPs (N=15) with topographies (middle row) and group averages (bottom row) shown at 8 peak times determined from the averaged source TEP (top row).

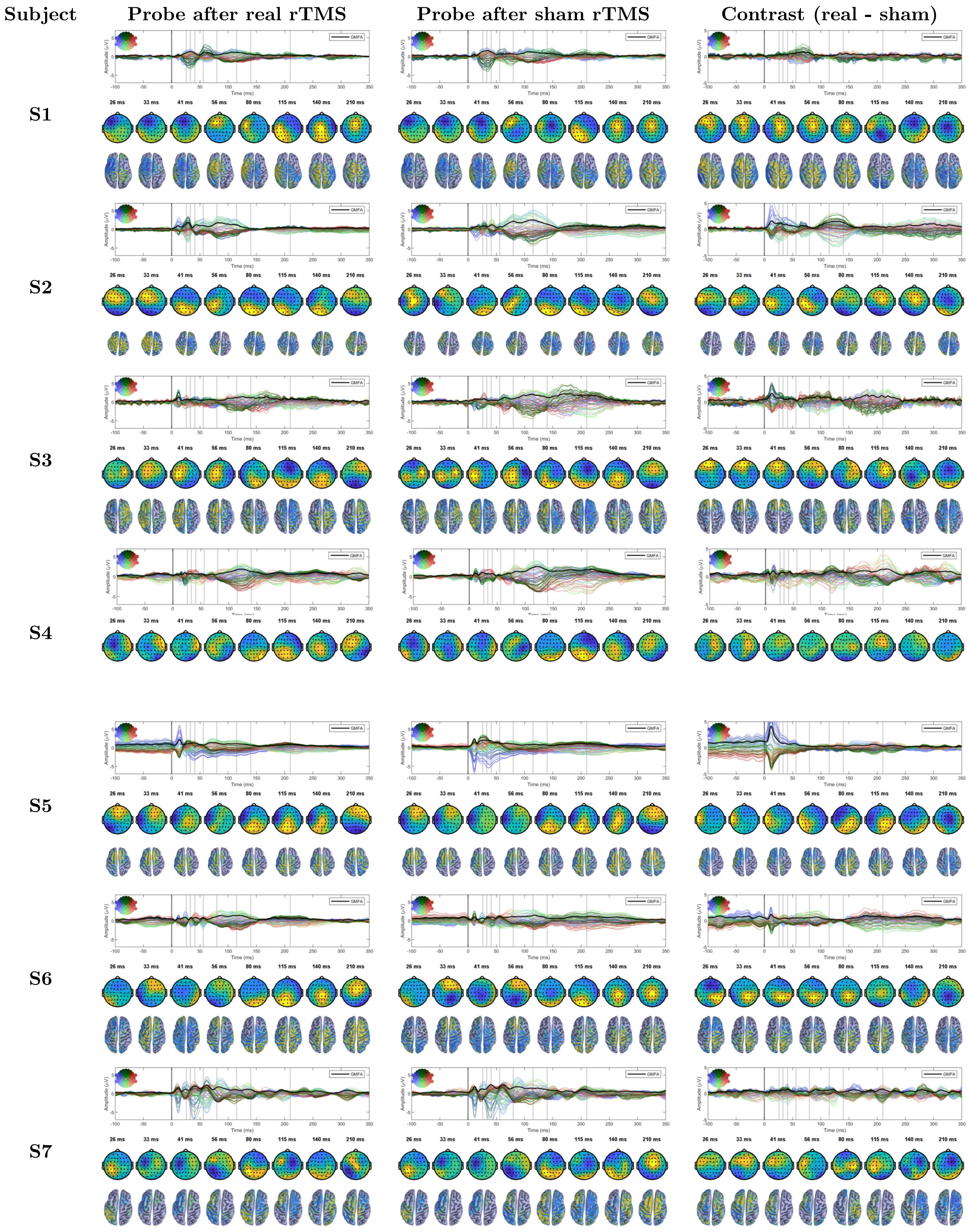

Figure S16 – continued on next page

Figure S16 – continued from previous page

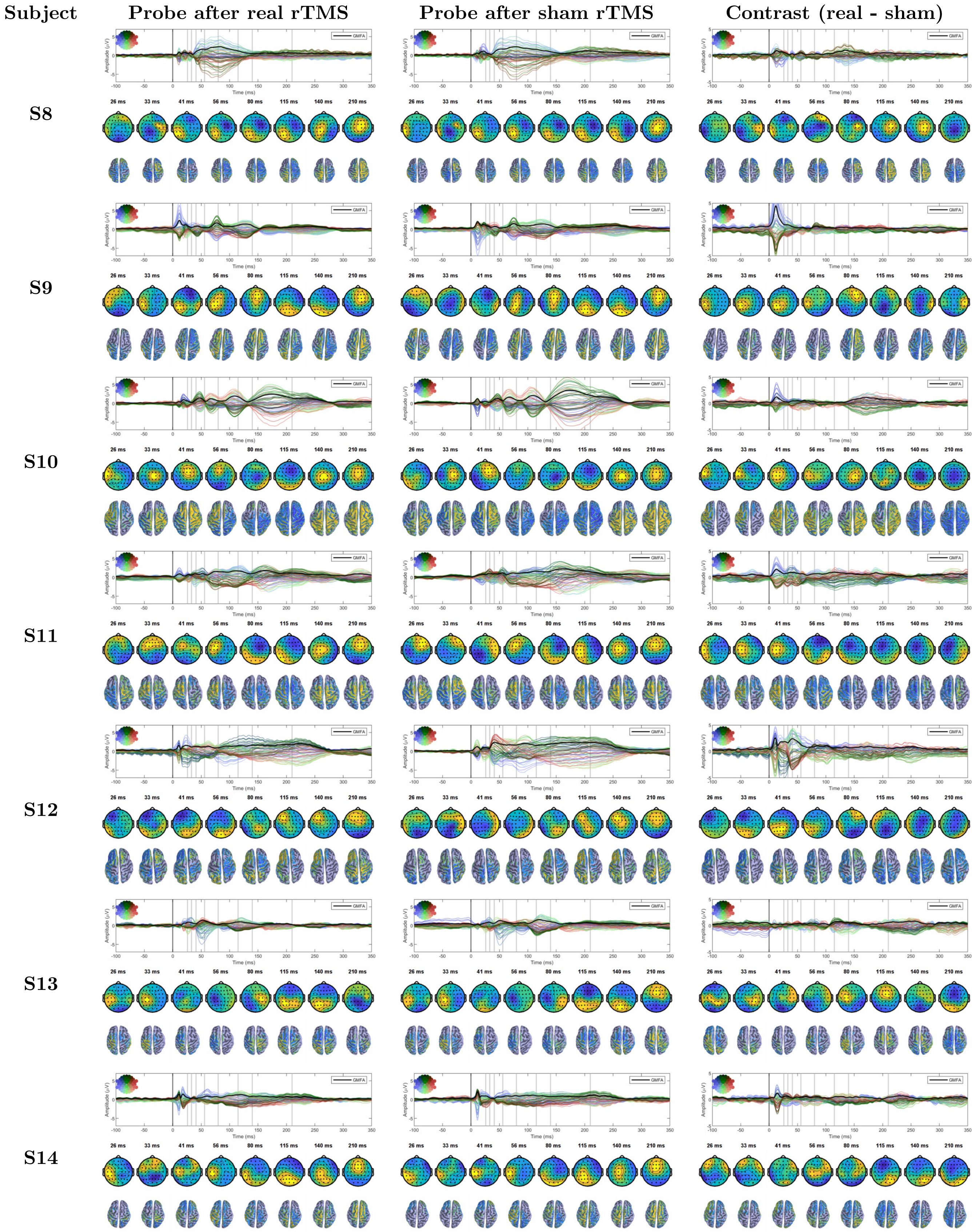

Figure S16 – continued on next page

| Subject | Probe after real rTMS | Probe after sham rTMS | Contrast (real - sham) |
|---|---|---|---|
| S15 | | | |
| S16 | | | |

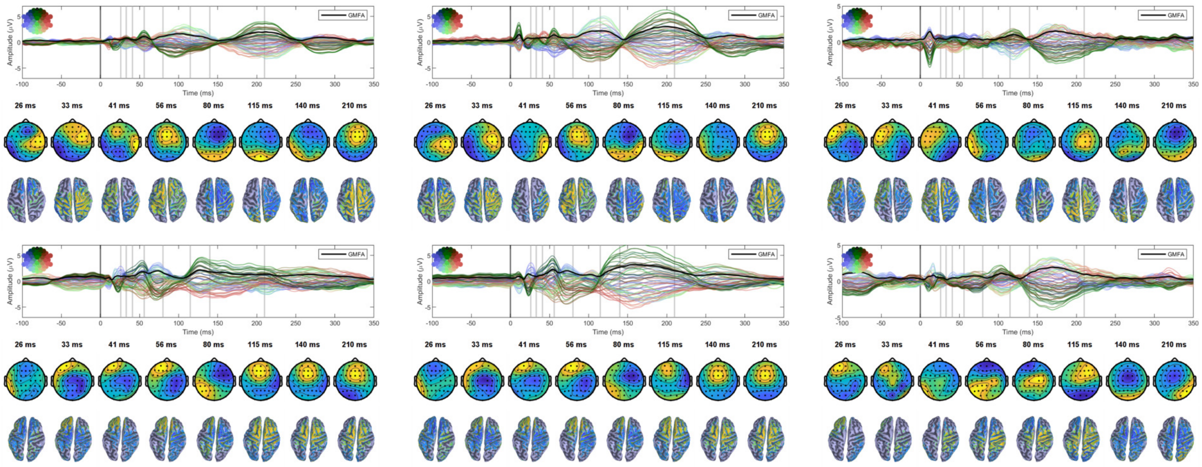

**Figure S16 | Single-subject butterfly plots and sensor and source space topographies.**
